# Early impairments of visually-driven neuronal ensemble dynamics in the rTg4510 tauopathy mouse model

**DOI:** 10.1101/2022.10.13.512017

**Authors:** Aleksandra Parka, Caroline Degel, Jakob Dreyer, Ulrike Richter, Benjamin Hall, Jesper F. Bastlund, Bettina Laursen, Maiken Nedergaard, Florence Sotty, Paolo Botta

**Affiliations:** Center for Translational Neuromedicine, University of Copenhagen, Blegdamsvej 3B, 2200 Copenhagen, Denmark; H. Lundbeck A/S, Neuroscience Research, 9 Ottiliavej, 2500 Valby, Denmark; H. Lundbeck A/S, Department of Bioinformatics, 9 Ottiliavej, 2500 Valby, Denmark

**Keywords:** Tauopathy, rTg4510, Alzheimer’s Disease, frontotemporal dementia, calcium imaging, pyramidal neurons, visual cortex, electroencephalography, diazepam, pentylenetetrazol

## Abstract

Tau protein pathology is a hallmark of many neurodegenerative diseases, including Alzheimer’s Disease or frontotemporal dementia. Synaptic dysfunction and abnormal visual evoked potentials have been reported in murine models of tauopathy, but little is known about the state of the network activity on a single neuronal level prior to brain atrophy. In the present study, oscillatory rhythms and single-cell calcium activity of primary visual cortex pyramidal neuron population were investigated in basal and light evoked states in the rTg4510 tauopathy mouse model prior to neurodegeneration. We found a decrease in their responsivity and overall activity which was insensitive to GABAergic modulation. Despite an enhancement of basal state coactivation of cortical pyramidal neurons, a loss of input-output synchronicity was observed. Spectral power analysis revealed a reduction of basal theta oscillations in rTg4510 mice. Enhanced susceptibility to a sub-convulsive dose of pentylenetetrazol was further indicated by an increase in theta power and higher number of absence-like seizures in rTg4510 compared to control mice. Our results unveil impairments in visual cortical pyramidal neuron processing and define aberrant oscillations as a biomarker candidate in early stages of neurodegenerative tauopathies.

## Introduction

Tau pathology-induced conditions are collectively labelled as tauopathies, where tau protein hyperphosphorylation and subsequent formation of neurofibrillary tangles (NFTs) lead to brain atrophy and memory impairment [1, 2]. Neurodegenerative diseases such as Alzheimer’s Disease (AD), frontotemporal dementia and cortico-basal degeneration are some of the classical examples of tauopathy [1]. Emerging evidence shows that tau oligomers, which are considered as precursors of NFTs, are more toxic to the healthy brain than tangle formations [3–6]. Additionally, while many studies explored the consequences of hyperphosphorylated tau in the late stages of pathology, little is known about the earlier stages where cognitive decline is not yet advanced but pre-tangle formations are already emerging in the brain [7–10].

Various animal models of tauopathy have been developed. Amongst those, the rTg4510 mouse model contains the P301L mutation under the calcium/calmodulin-dependent kinase II alpha (CaMKIIα) promoter restricting overexpression of human tau (h-tau) to the excitatory pyramidal neurons (PNs) of the forebrain [9,11,12]. This mouse model shows progressive neurodegeneration from approximately 6 months of age [10, 13]. Despite the lack of major morphological changes in PNs, disparate findings showing excitability impairments, synaptic dysfunctions, and network and oscillatory imbalance were reported at early stages of tau pathology [10,14–19]. Not only intrinsic membrane properties of PNs and their neuronal activity were found to be altered in *ex vivo* and *in vivo* studies [14, 20], but also dysfunctional inhibitory neurotransmission in young rTg4510 mice [15,19,21] . As a result, multifactorial impairments of the intrinsic and extrinsic neuronal properties reflected in single-cell excitability, cortical long-range and local interconnectivity might lead to altered PNs activity and cortical processing prior to neurodegeneration [22–24]. Collectively, these alterations could affect cortical oscillatory activity which is otherwise associated with healthy brain function [25–27]. In fact, neural oscillations show a shift of power spectra towards lower frequencies in prodromal stages of AD and rodent tauopathy models [20,28–30]. Aberrant oscillatory activity in AD might be associated to the reported higher incidence of epileptiform activity [31–33]. In addition, tau pathology was found in epilepsy patients and elimination or reduction of tau prevented seizures and network hyper-synchrony in tauopathy mouse models [34–36].

We have previously discovered an early tauopathy biomarker in the visual evoked potentials (VEPs) recorded from primary visual cortex (V1) of young rTg4510 mice prior to neurodegeneration, which was positively correlated with levels of hyperphosphorylated tau resembling pre-tangle formations [37]. Moreover, as PNs constitute the main neuronal population in cortical layers, we hypothesized that impairment of their physiological activity induced by tau pathology may result in overall abnormal network processing during visual stimulation [27, 38]. Based on abnormal neuronal processing found in separate studies [14,20,29,37], dysfunction of PNs in the visual cortex (^V1^PNs) could explain aberrant VEPs found in rTg4510 mice in early stages of pathology preceding neuronal loss [37]. To our knowledge, state dependent ^V1^PNs activity in rTg4510 mice prior to neurodegeneration has never been studied on a single cell level *in vivo*. In the present study, we investigated the consequence of tau overexpression in young freely moving rTg4510 mice on cortical visual processing using single-cell calcium imaging from thousands of defined ^V1^PNs during visual stimulation and their response to GABAergic tone modulation. Additionally, we explored whether electroencephalogram (EEG) oscillations in the visual cortex in response to a sub-convulsive dose of pentylenetetrazole (PTZ) could be related to single-cell dynamics, and therefore potentially represent an early-stage biomarker for tauopathies.

## Materials and methods

### Animals

Age-matched, 3-4 months old male mice were bred at Taconic, Denmark. rTg4510 animals are a result of a cross between a tetracycline-controlled transactivator protein (tTA) allele under a CaMKIIα promoter and an allele containing a P301L mutation on the tau gene (MAPT) and a tetracycline-responsive promoter element (TRE) [9]. Mice expressing tTA were maintained on the 129S6 background strain and mutant tau_P301L_ responder mice were maintained on the FVB/NCrl background strain. As there are sex-dependent differences in rTg4510, male mice were exclusively used in this study [39]. Upon arrival they were single housed with *ad libitum* access to food and water. The light/dark cycle was maintained at 12 h with lights on at 6 am. All animal experiments were performed in accordance with the European Communities Council Directive #86/609, the directives of the Danish National Committee on Animal Research Ethics and Danish legislation on experimental animals. Age-matched tTA animals were used as controls in this study and will be referred to as controls throughout.

### Study design

Calcium imaging was performed in 14 rTg4510 and 15 control animals. Vehicle, diazepam and PTZ treatments were performed in a pseudo-random crossover design, where every animal was dosed with each compound with a 7-day wash-out period. EEG recordings performed before and during PTZ treatment were performed in a separate cohort of animals consisting of 16 rTg4510 and 20 control mice.

### Pharmacological treatments

Diazepam, which is a positive allosteric modulator of the GABA_A_ receptor, was dosed at 1 mg/kg s.c. and PTZ, a GABA_A_ receptor antagonist, was administered at 35 mg/kg i.p.. Both compounds were dissolved in sterile saline which was also administered in vehicle treated animals and injected at 10 ml/kg. Since PTZ at 35 mg/kg did not trigger behavioural seizures in our pilot study, it was considered as a sub-convulsive dose (Table S1). Special care was taken to note any seizure-like behavioural manifestations during the imaging session via the cameras placed in the recording arena. All pharmacological compounds were synthesized at Lundbeck (H. Lundbeck, Valby, Denmark).

### Stereotaxic surgery for viral injection and lens implantation

The surgery area was sanitized, and isoflurane anaesthesia was induced at 5 vol% and an airflow of 0.3 and 0.7 L/min of O_2_ and N_2_O respectively. Anaesthesia was maintained by decreasing the flow of isoflurane to approximately 2 vol%. The head was shaved and disinfected with chlorhexidine followed by a subcutaneous (s.c.) injection of Marcain (2.5 mg/ml bupivacaine, AstraZeneca, Albertslund, Denmark). The incision was made through the middle and skull was wiped down and dried off. Animals were mounted in a stereotaxic frame (David Kopf Instruments, Tujunga, CA, USA) and the coordinates of bregma and lambda were identified. Adjustments were done if the head was not levelled, i.e., if bregma and lambda were not aligned in terms of depth and lateral coordinates. A cranial window (1 mm x 1 mm) was made, and the animals were unilaterally injected with 400 nL of ssAAV-9/2-mCaMKIIα-jGCaMP7f-WPRE-bGHp(A) (Neuroscience Center Zurich, Zurich, Switzerland; titer: 1.1×10^13^ vg/mL) in the primary V1 area (AP (anteroposterior): −2.92, ML (mediolateral): −2.2). To ensure homogeneous viral expression, 10 nL was released in 10 pulses at four depths (DV (dorsoventral): −1.35, −1.1, −0.8 and −0.5), starting at the deepest coordinate and retracting the tip of the capillary by 0.3 um after every 100 µL deposit. The injection process was automatized by using Nanojet III Injector (Drummond Scientific, USA). To place the lens implant, an incision was made at the edge of the cranial window in which the prism lens (ProView™ Integrated Prism Lens 1.0 mm x 4.3 mm: 1-mm-diameter gradient index (GRIN) lens, length 4.3 mm, 1050-004419; Inscopix, Palo Alto, CA, USA) was gently lowered until reaching the predetermined coordinates (AP: −1.35, ML: −2.2 DV: −1.35). The exposed brain was covered with Kwik-Cast silicone sealant (World Precision Instruments, Friedberg, Germany) and the protruding lens and base of the implant was covered with C&B Metabond cement (Parkell, Long Island, NY, USA). As post-surgery care, the mice were weighed and treated with Noromox (150 mg/ml amoxycilin hydrate, ScanVet, Fredernsborg, Denmark) and Norodyl (50 mg/ml carpofen, ScanVet, Fredensborg, Denmark) for 5 days.

### Calcium imaging in freely moving animals

Prior to commencing the calcium imaging experiments, each animal was tested after a minimum of three weeks from viral injection to ensure that fluorescent signal could be acquired from single cells. Animals with no neurons detected were excluded from the study. Each recording arena consisted of a transparent cylinder 20 cm in diameter with a strip of light-emitting diode (LED) lights wrapped around it, 5 cm above the base. All animals were habituated on three consecutive days by connecting them to a dummy miniscope (1050-003762; Inscopix, Palo Alto, CA, USA) and placing them in an arena for 1 h, with an intraperitoneal (i.p.) saline injection after 30 min. On the day of the imaging, the animal was connected to the miniscope (excitation: blue LED; excitation filter: 475/10 nm, 0.24-0.6 mW/mm2; emission filter: 535/50 nm; Inscopix, Palo Alto, CA, USA) and placed in the experimental arena. The video was acquired by the Inscopix software (Inscopix, Palo Alto, CA, USA) at 20 Hz. Additionally, the intensity of LED light in the microscope, focal plane and gain parameters were set to be optimal for each individual animal prior to the start of each recording session.

### Light stimulation paradigm in calcium imaging

The animals were subjected to a light stimulation protocol in which a 50-ms single white light stimulus was presented at 5, 10, 25, 50 and 100% (5, 10, 460, 2540 and 6390 lux at the centre of the arena, respectively) intensity of the LED strip. The stimulus intensities were shown on a randomized basis with 25-35 s between flashes. Images were acquired 10 s prior to and proceeded for 10 s after stimulus onset. The baseline consisted of 25 flashes (each intensity displayed five times) after which the drug or vehicle was administered. The recording was paused for 10 min post-administration and resumed with the same paradigm as in the baseline.

### Miniscope imaging analysis

Raw imaging videos were analysed using the Python API from the Inscopix Data Processing Software release 1.3.1 (IDPS, Inscopix, Palo Alto, CA, USA), followed by a quality control check step. Using the API, the data was pre-processed by spatial binning of 4×4 pixels and motion correction. The API implementation of the algorithm for constrained non-negative matrix factorization for endoscopic data (CNMF-e) [40, 41]. The subsequent quality control was based on two characteristics of putative cells: either the shape was considered unlikely for a well-isolated single PN; or the predicted calcium timeseries for the single putative cell appeared non-stereotypical, with long, smooth segments, sometimes even constant, lacking the natural complexity and randomness typically present in calcium data.

In more detail, the shape test was based on calculating 3 parameters of the shape of the putative cell: area, radius of gyration, and the ratio of the two principal axes of the cell shape. The minimum covariance determinant (MCD) method was then used to detect outliers in this multivariate data [42]. In practice, we discarded putative cells exceeding an upper limit to the robustly calculated Mahalanobis distance. The MCD method was implemented in Python using MinCovDet from scikit-learn package (version 1.1.2).

As a test of the calcium signal from putative cells, we calculated the Permutation entropy (PE) of the first derivative of the time series [43]. Here we discarded putative cells where the PE of the calcium traces did not meet a lower limit. The PE was calculated in Python using the pyentrp package (version 0.7.1). Altogether, we found that around 10% of the cells detected by the CNMF-e algorithm were outliers and should be excluded from the analysis (Fig. S1).

Responsivity and response parameters of ^V1^PNs were calculated by calculating a z-score. Individual z-scores from each neuronal trace were calculated by comparing raw activity to calcium activity during 5 s baseline prior to light stimulation:

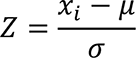

Where x_i_ represents raw calcium value for each frame of the trace, µ is the baseline average and σ indicates the baseline standard deviation. The z-score threshold was set at a range between −1.96 and 1.96 (significance of p<0.05, 95% confidence), where if 5 consecutive frames (corresponding to 250 ms) exceeded the upper or lower threshold, the neuron was termed excited or inhibited, respectively, while neurons within the threshold limits were considered to be silent upon light stimulation.

Response latency was considered as the first time point at which the trace reached the upper or lower z-score threshold, whereas the response duration was the time period between the time points of initial rise above and consequent drop below the z-score threshold for putatively excited cells, and vice versa for putatively inhibited cells. Finally, the magnitude of response was established by calculating the area under the curve (AUC) of the trace surpassing the z-score threshold.

Furthermore, the Online Active Set method to Infer Spikes (OASIS) algorithm as described by Friedrich and colleagues [44] was applied to the z-scored calcium traces to extract calcium events in imaged cells. More specifically, spikes surpassing a signal-to-noise ratio of 3 were defined to determine a ^V1^PN event. To investigate the connectivity of detected ^V1^PNs, the Pearson cross-correlation coefficient (CC) was calculated at lag 0 for all possible pairs of neurons in MATLAB (MathWorks, Natick, Massachusetts, USA) as follows:

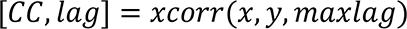

where x and y represent the z-scored traces of one possible pair of ^V1^PNs and maxlag is equal to 10 s. The final CC for each neuron was an average of the CC values over all its possible ^V1^PN pairs. CC = 1 were not included as they represent comparison amongst the same neurons. In order to evaluate ^V1^PN CC change over time, CC lag ranged between –10 to 10 s from stimulation.

### Tissue extraction and lens placement validation

Mice were injected with an overdose of Avertin and perfused with heparinized KPBS. The brains were kept in 4% paraformaldehyde for 48 h and then in 0.01% sodium azide and KPBS solution. The brains were freeze-sectioned at 35 µm in the coronal plane and mounted on slides with ProLong™ Diamond Antifade Mountant with DAPI (Thermo Fisher Scientific). Sections were visualized using confocal microscopy (Leica TCS SP8, Leica, Wetzlar, Germany) with x40 (HC PL APO 40/0.85, Leica, Wetzlar, Germany) to validate the expression of the virus as well as the localization of the implant. Viral expression was quantified by measuring intensity of staining in cortical layers I-VI in ImageJ. After assessing the limits of the lens implant, the middle image was taken.

### Stereotaxic surgery for placement of EEG electrodes

Surgery was carried out as described by Parka et al. [37]. Briefly, mice were anesthetized with isoflurane (30% oxygen/70% nitrogen; 5% isoflurane for induction, 1.5-2% for maintenance) and treated with Noromox (150 mg/ml amoxycilin hydrate, ScanVet, Fredernsborg, Denmark) and Norodyl (50 mg/ml carpofen, ScanVet, Fredensborg, Denmark). Marcain (2.5 mg/ml bupivacaine, AstraZeneca, Albertslund, Denmark) was injected s.c. to the skull area and vaseline-based ointment was applied on the eyes. Screw electrodes (E363/20/1.6/SPC, PlasticsOne, Roanoke, VA, USA) were inserted in the following coordinates: one electrode over the visual cortex (AP: −3.6 mm, ML: 2.3 mm), two electrodes in the frontal cortex (AP: +2.8, mm ML: +/-0.5 mm), one electrode in parietal cortex (AP: −2 mm, ML: 2 mm), one ground electrode (AP: −2 mm, ML: −2 mm) and one reference electrode in the most posterior part of the skull (AP: −6 mm, ML: −2 mm). Only the signal from visual cortex electrode was analyzed while the rest of the electrodes served a stabilizing purpose. Animals were treated with Norodyl and Noromox for 5 days post-surgery and full recovery lasted 10 days.

### EEG recordings in freely moving animals

Recovered animals were submitted for EEG recordings during the dark phase after 2 weeks of habituation for approximately 3 h once a week. The first hour of recording was considered as baseline EEG activity. VEP stimulation commenced right after the recording of baseline. Afterwards, the mice were recorded immediately after application of PTZ. The signal was sampled at 1 kHz (CED Power 1401, Cambridge Electronic Design Ltd, Cambridge, UK), passed through a band-pass filter of 0.01-300 Hz (Precision Model 440; Brownlee, Palo Alto, CA, USA) and amplified 1000 times. EEG waveforms were acquired with Spike2 version 7.2 (Cambridge Electronic Design Ltd, Cambridge, UK) and analyzed with MATLAB software (MathWorks, Natick, Massachusetts, USA). VEP recordings were performed as described by Parka et. al [37], white light stimuli lasted 10 ms with an inter-stimulus interval of 3 s for 5 and 21.5 lux (150 repeats), 4 s for 45 lux (150 repeats), 5 s for 73.5 and 108 lux (both 100 repeats), 7 s for 140 lux (75 repeats) and 10 s for 177 lux (50 repeats).

### EEG recording analysis

EEG recording waveforms were analysed in Spike2 version 7.2 (Cambridge Electronic Design Ltd, Cambridge, UK). For the power spectral analysis, the signal was passed through fast Fourier transformation using a Hanning window resulting in a frequency resolution of 0.49 Hz. Frequency bands delta (δ), theta (θ), alpha (α) and beta (β) were defined by 1-4, 4-10, 10-14 and 14-20 Hz, respectively. The power of each frequency band was normalized to the total power in the range 1-100 Hz. The peak θ frequency in each animal was defined as the frequency at which the highest power was detected in the θ frequency band. EEG recordings obtained from animals after PTZ administration were band passed through a 4-8 Hz Butterworth 3^rd^ order filter. Epileptiform activity was defined as absence seizure-like episodes when events had amplitudes surpassing pre-PTZ baseline max range and exceeded 1s in duration. Quantification analysis was done by manual inspection of filtered traces by an experimenter.

### Statistics

All statistics were performed in GraphPad Prism (GraphPad Software, San Diego, California, USA). For comparison between multiple groups with one factor, one-way ANOVA was employed. When comparing differences between multiple groups with two factors, a two-way ANOVA test was performed followed by Šidák’s multiple comparison post hoc analysis provided that one of the factors had significant effect. The two-tailed student t-test was used when comparing means of two groups. If p<0.05 the difference in comparison was considered significant and precise p-values are stated in the results section, while p>0.05 was considered to indicate a non-significant difference and no further indication is given.

## Results

### Bulk fluorescence response to the light stimulation is decreased in rTg4510 mice

As visual evoked potentials recorded in V1 were altered at early stages of tauopathy [37], we tested whether cortical processing was impaired in young rTg4510 compared to controls at the neuronal level. Neuronal activity of ^V1^PNs was imaged from both genotypes four weeks after injecting an AAV virus expressing the calcium sensitive GCaMP7f biosensor and implanting a prism lens along the dorso-ventral axis of V1 (Fig. 1A, S2). Quantitative post-hoc imaging analysis showed that viral expression and calcium imaging plane centre was largely restricted to the deep layers of V1 in both control and rTg4510 animals (Fig. 1B, S2). The flash of light elicited a marked bulk response from the imaged area (Fig. 1C, D) that significantly increased with light intensity in both control and rTg4510 animals (p=0.0053 and 0.0005 for control and rTg4510 mice, respectively, by one-way ANOVA, intensity effect, Fig. 1E, F). However, the peak response imaged from rTg4510 mice was significantly lower than the response elicited in control animals (p=0.0193 by two-way ANOVA, genotype effect, Fig. 1F).

**Figure 1.**
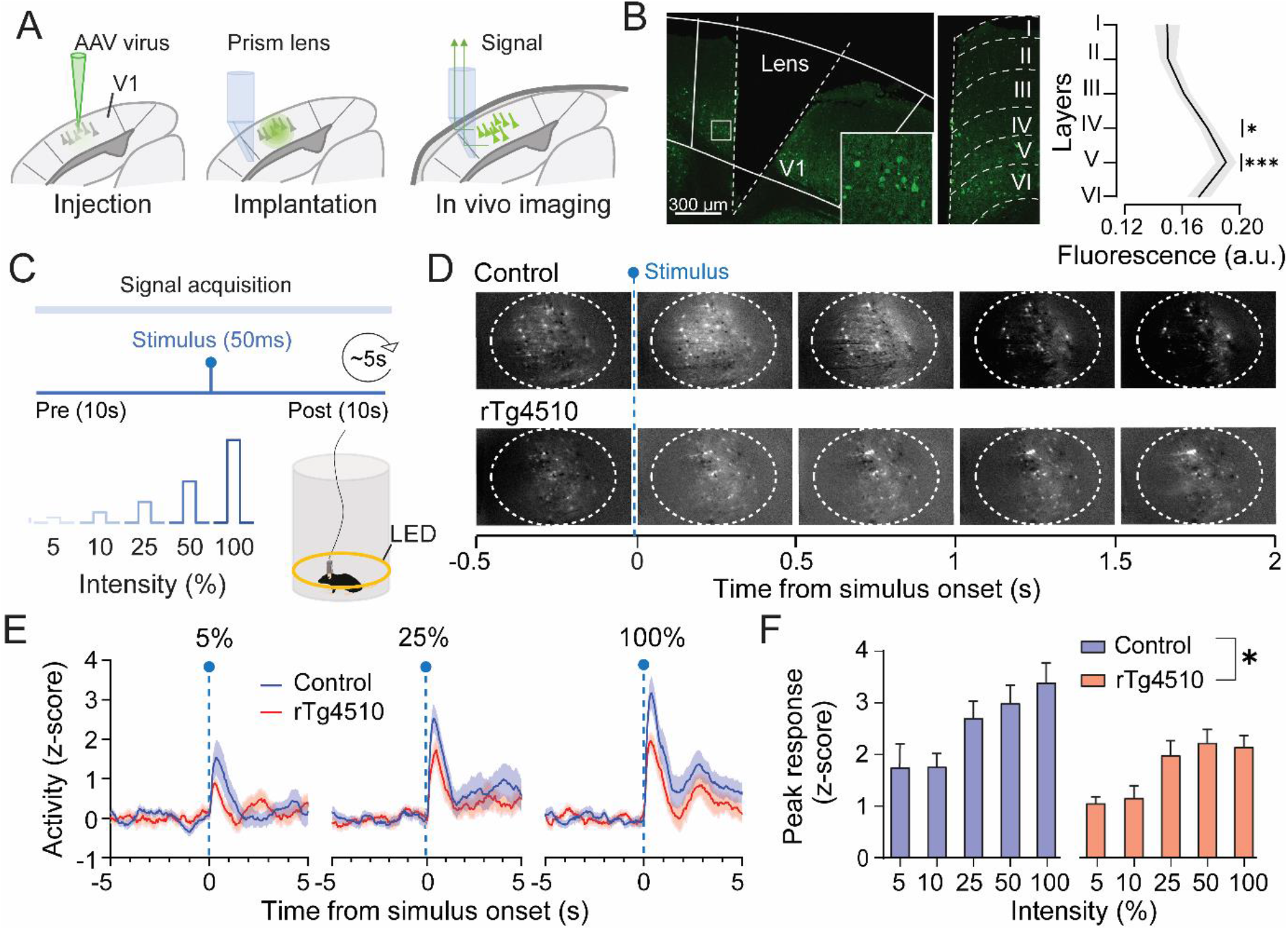
Altered ^V1^PNs bulk response to light stimulation in the rTg4510 mouse model. (A) Schematics of AAV-CaMKII-GCamP7f viral injection and prism lens implantation followed by imaging procedure. (B) Representative confocal image of lens placement in the visual cortex with a magnified image of labelled ^V1^PNs (left, square with white borders). Image of the focal plane across cortical layers (centre) and bar graph of the viral expression across cortical layers (right, n=17) (*) p<0.05, (***) p<0.001 by one-way ANOVA, layer effect. Gray area around the line represents SEM. (C) Light stimulation protocol and a scheme of the imaging arena with a freely moving animal connected to the imaging system. (D) Representative time course of light induced neuronal activation of ^V1^PNs in control and rTg4510 animals. Stimulus occurred at time point 0 and is marked with a dashed blue line. ROI for bulk image analysis is shown with a white circle. (E) Time course of light evoked bulk activity (z-scores) in control (blue, n=15) and rTg4510 mice (red, n=14). (F) Bar graph of peak evoked responses (z-scores) of ^V1^PNs responses in control (blue, n=15) and rTg4510 mice (red, n=14) as a function of light intensity. (*) p<0.05 by two-way ANOVA, genotype effect. Data are shown as mean ± SEM.

### Light responsivity of individual ^V1^PNs is decreased in rTg4510 mice

Based on the changes showed using bulk imaging, we investigated if ^V1^PN evoked activity was altered in rTg4510 mice at single cell level (Fig. 2A). As the total number of neurons detected with CNMF-e algorithm was not significantly different between control and rTg4510 animals, we performed the analysis using approximately the same size neuronal population (Fig. 2B). Neuronal subtypes were selected based on their post-stimulus response and sorted into three subclasses: excited (∼5-20%), silent (∼70-90%), and inhibited neurons (∼2-8%; Fig. 2C-F, S3). We found that the fraction of excited neurons significantly increased with the intensity of the light in control and rTg4510 mice (p<0.0001 and p=0.0004 in control and rTg4510, respectively, by one-way ANOVA, intensity effect). However, the fraction of excited neurons was significantly lower in rTg4510 animals compared to controls (p=0.0129, by two-way ANOVA, genotype effect; Fig. 2E). Similarly to the excited ^V1^PNs, increasing light intensity enhanced the inhibited population fraction (p<0.0001, 0.036 in control and rTg4510, respectively, by one-way ANOVA, intensity effect) although to a lesser extent in rTg4510 mice compared to controls (p=0.0184, by two-way ANOVA, genotype effect; Fig. S3). As expected, the fraction of silent neurons decreased with increasing light intensity in both genotypes (p<0.0001, 0.0005 for control and rTg4510 mice, respectively, by one-way ANOVA, intensity effect; Fig. 2F) and to a larger extent in rTg4510 compared to control mice (p=0.0064 by two-way ANOVA, genotype effect; Fig. 2F). Interestingly, none of the response parameters of excited nor inhibited neurons such as magnitude, duration or latency were consistently altered between control and rTg4510 animals (Fig. 2G-I, S3).

**Figure 2.**
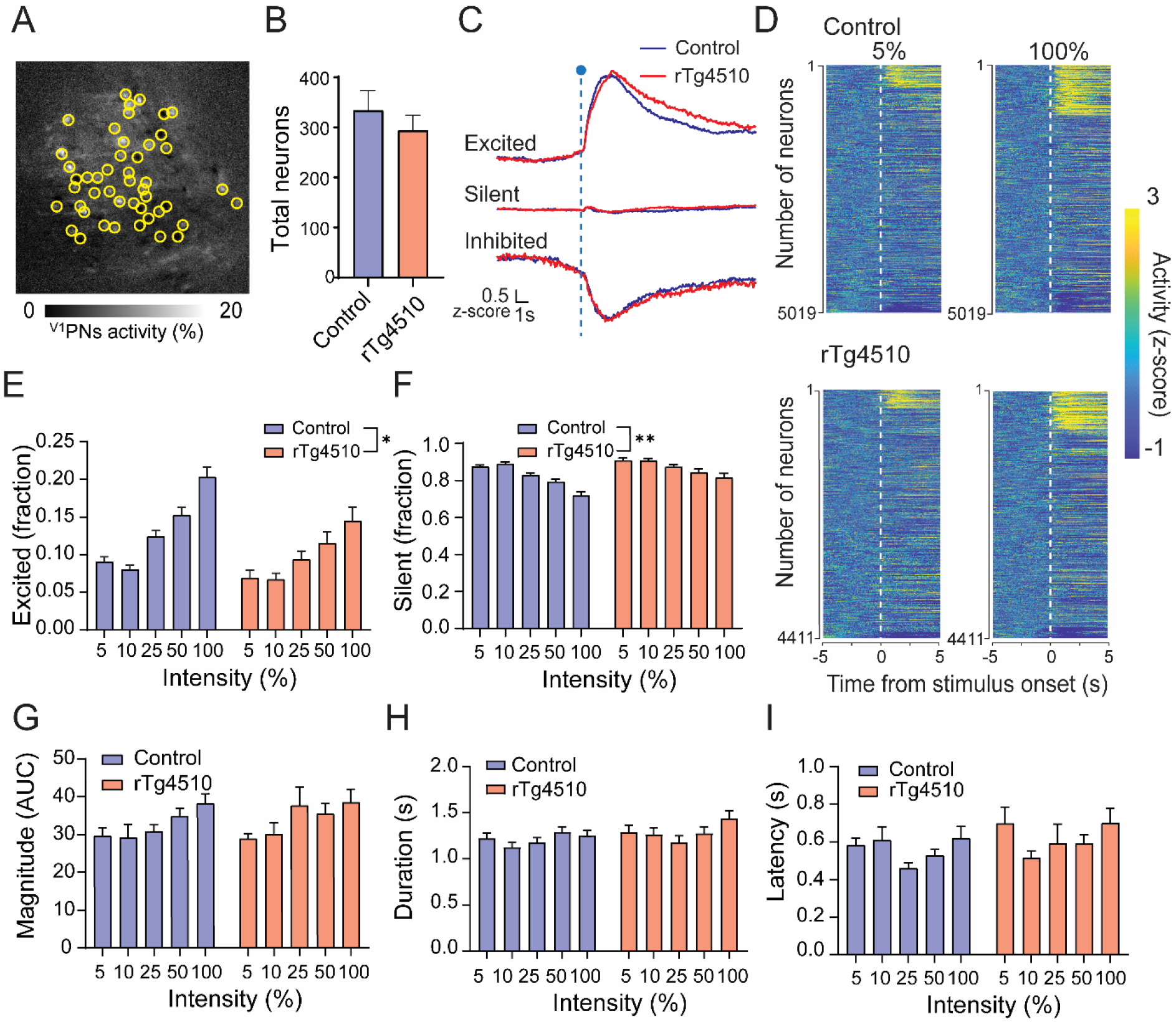
Impairment in responsiveness of ^V1^PNs in the rTg4510 mouse model. (A) CNMF-E-processed video frame of the V1PNs. Neurons passing the quality control standards are selected in yellow. (B) Average number of neurons detected from control (blue, n=15) and rTg4510 (red, n=15) animals. (C) Average activity traces of ^V1^PNs in control (blue, n=1004, 3617, 400 neurons for excited, silent and inhibited, respectively) and rTg4510 (red, n=691, 3522, 199 neurons for excited, silent and inhibited, respectively) evoked by 100% intensity of light. (D) Activity of all ^V1^PNs evoked by 5 and 100% of light in control and rTg4510 animals. Stimulus occurred at time point 0 and is marked with a white dashed line. (E) Excited and (F) silent fractions of all detected neurons in control (blue, n=15) and rTg4510 mice (red, n=15). (*) p<0.05, (**) p<0.01 by two-way ANOVA, genotype effect. (G) Magnitude, (H) duration and (I) latency of responses in excited ^V1^PNs in control (blue, n=15) and rTg4510 animals (red, n=15). Data are shown as mean ± SEM.

### GABAergic modulation of neuronal activity is attenuated in rTg4510 mice

As impaired GABAergic transmission has been reported in the rTg4510 mouse model [15], we tested whether the modulation of ^V1^PN excitability in the evoked state by a low dose of either diazepam or PTZ was altered in rTg4510 mice (Fig. 3A). We found that diazepam significantly decreased the fraction of excited neurons at all intensities in control (p=0.0108, 0.007, 0.0105 for 5%, 25% and 100%, respectively, by two-way ANOVA, treatment effect) but not in rTg4510 mice (Fig. 3B). Diazepam induced an increase in silent fraction of ^V1^PNs in control mice (p=0.0072, 0.0039, 0.0227 for 5%, 25% and 100%, respectively, by two-way ANOVA, treatment effect, Fig. 3C) but not in rTg4510 mice. As the fraction of inhibited ^V1^PNs amounted for only 2-8% of total detected neurons, it was not further analysed. On the other hand, treatment with PTZ did not induce any significant change in responsivity of ^V1^PNs in control or rTg4510 animals (Fig. 3B).

**Figure 3.**
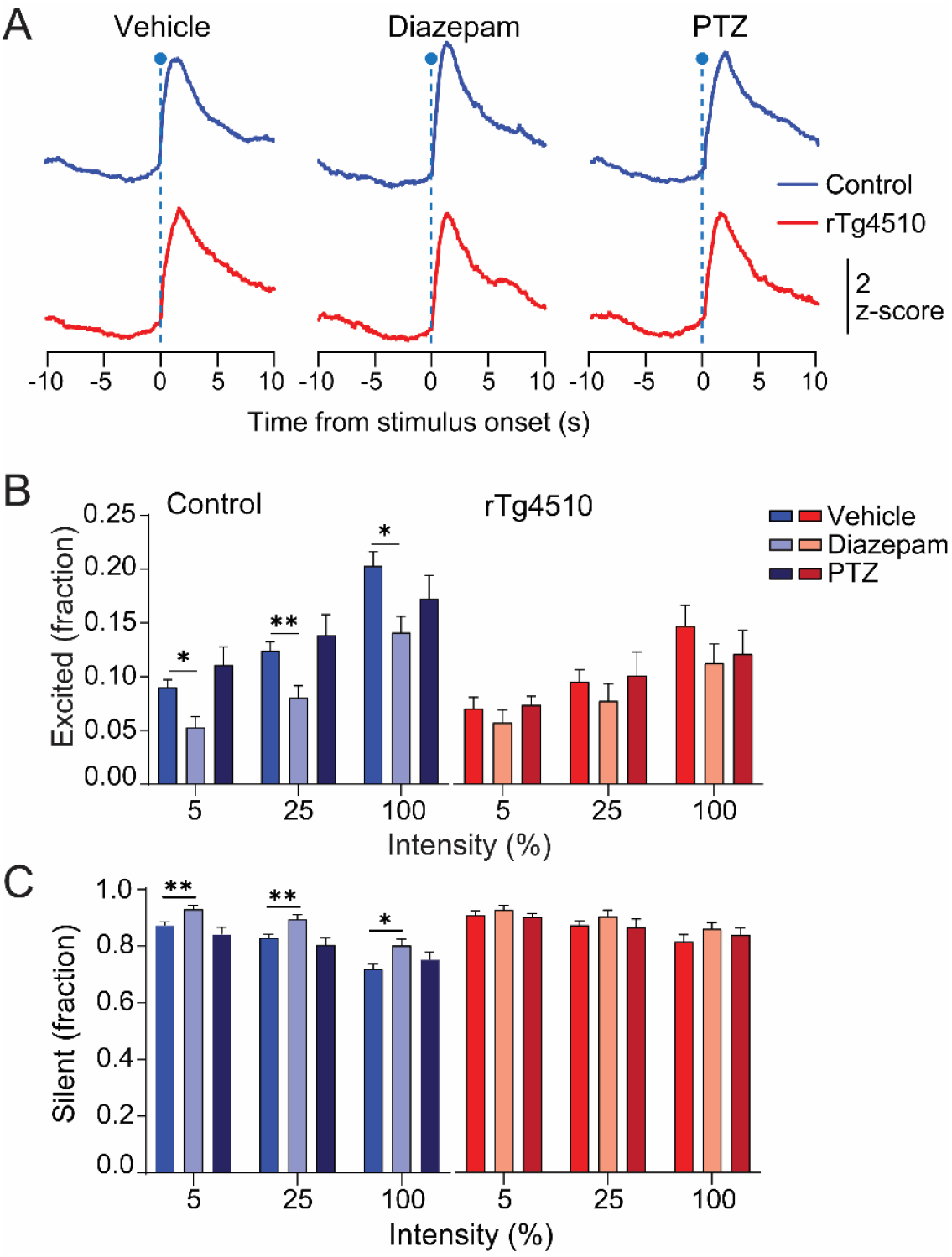
Differential effect of pharmacological GABAergic treatments on ^V1^PN responsivity. (A) Average calcium traces of ^V1^PN activity evoked by 100% of light intensity under the influence of different pharmacological treatments in control (blue, n=1004, 561, 760 neurons for vehicle, diazepam and PTZ respectively) and rTg4510 animals (red, n=691, 521, 508 neurons for vehicle, diazepam and PTZ respectively). Dashed blue line represents the stimulus onset. (B) Excited and (C) silent ^V1^PNs fractions at incremental light intensities in control (blue; n = 15, 12, 13 mice for vehicle, diazepam and PTZ, respectively) and rTg4510 mice (red; n = 15, 16, 15 mice for vehicle, diazepam and PTZ, respectively). (*) p<0.05, (**) p<0.01 by two-way ANOVA, drug effect. Data are shown as mean ± SEM.

### Basal activity of ^V1^PNs is attenuated and unresponsive to GABAergic modulation in rTg4510 mice

To investigate basal activity of ^V1^PNs, single cell pre-stimulus neuronal events were isolated and analysed in baseline and under treatment conditions (vehicle, diazepam and PTZ; Fig. 4A, B). Analysis of pre-stimulus event rates in baseline of single ^V1^PNs showed a significant decrease in rTg4510 animals compared to controls (p=0.0001 by student t-test; Fig. 4B, C). While pre-stimulus event rates in control mice were significantly decreased by diazepam (p<0.0001 by two-way ANOVA, treatment effect) and increased by PTZ (p<0.0001 by two-way ANOVA, treatment effect), none of the compounds induced significant change in event rates in rTg4510 mice (Fig. 4D). Thus, the effects of both diazepam and PTZ were significantly different in rTg4510 compared to control mice (p<0.0001 by one-way ANOVA for both diazepam and PTZ, Fig. 4D).

**Figure 4.**
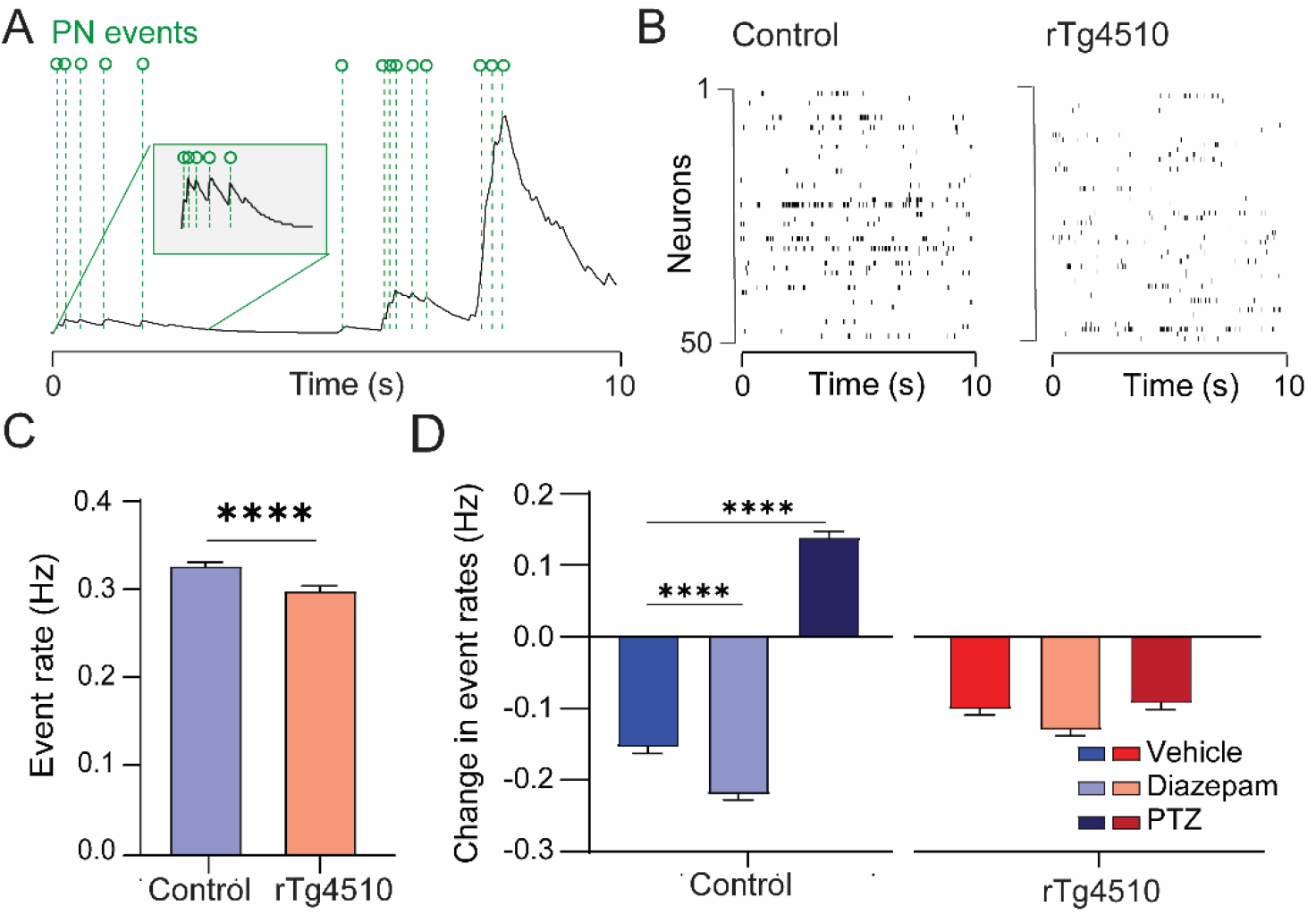
GABAergic modulation of basal activity of ^V1^PNs is altered in the rTg4510 mouse model. (A) Schematic of event deconvolution detection. ^V1^PN events are labelled with green dashed lines. (B) Deconvolved pre-stimulus events in a sample of 50 neurons in control and rTg4510 mice. (C) Average event rate in control (blue, n = 5019 neurons) and rTg4510 animals (red, n = 4411 neurons). (***) p<0.001 by student-t-test. (D) Average event rates under drug conditions in control (blue; n = 14, 13, 13 mice for vehicle, diazepam and PTZ, respectively) and rTg4510 mice (red; n = 15, 16, 15 mice for vehicle, diazepam and PTZ, respectively). (***) p<0.001, (****) p<0.0001 by two-way ANOVA, genotype and treatment effects. Data are shown as mean ± SEM.

### State-dependent network synchronicity is dysregulated in rTg4510

We investigated if synchronicity of the neuronal activity was affected at early stages of tauopathy by performing cross-correlation (CC) analysis of ^V1^PN pairs in basal and evoked state in the rTg4510 mice (Fig. 5A). First, we tested whether light stimulation enhanced network synchronicity by comparing the CC of excited ^V1^PN pairs between the basal and evoked states. As expected, we found that this measure increased in both genotypes (p<0.0001 for both genotypes in basal vs. 5% light; p<0.0001 for both genotypes in basal vs. 100% light by two-way ANOVA; Fig. 5B, C).

**Figure 5.**
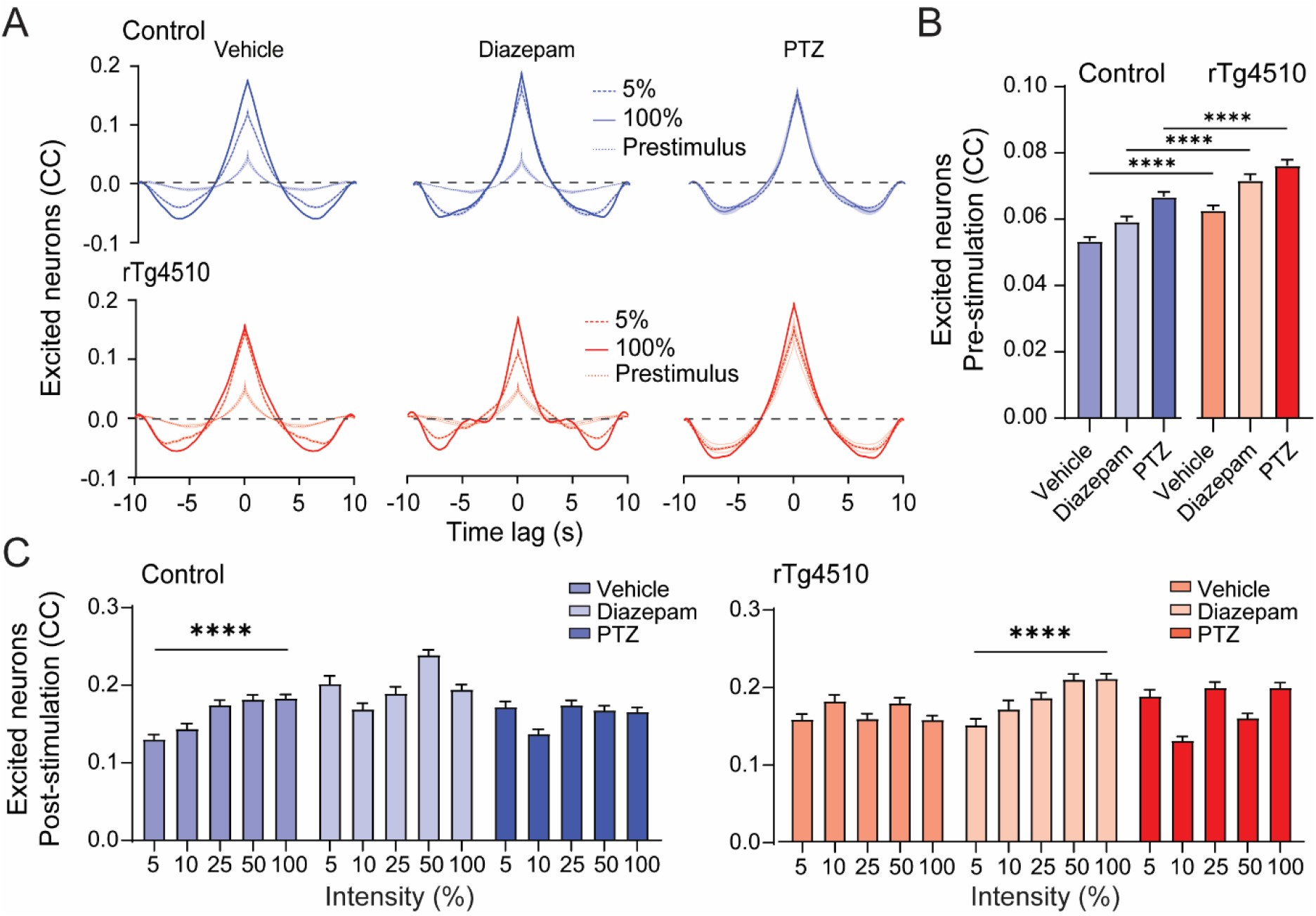
Enhanced cross-correlation between pairs of ^V1^PNs but input-output function impairment in rTg4510 mice. (A) Cross-correlation (CC) between pairs of ^V1^PN population at different lag times in 5 and 100% intensity as well as pre-stimulus period in control (top, blue, n=13, 13, 12 mice for vehicle, diazepam and PTZ, respectively) and rTg4510 mice (bottom, red, n=13, 14, 14 mice for vehicle, diazepam and PTZ, respectively). Shaded areas around lines represent standard deviation. (B) Pre-stimulus CC of excited (top) and silent (bottom) ^V1^PNs in control (n=2847, 1692, 2609 ^V1^PNs for vehicle, diazepam and PTZ, respectively) and rTg4510 mice (n=2212, 1809, 2241 ^V1^PNs for vehicle, diazepam and PTZ, respectively). (****) p<0.0001 by two-way ANOVA, genotype effect. (C) Post-stimulation cross-correlation coefficients in controls (left; n = 389, 364, 555, 654, 885 ^V1^PNs for vehicle; n = 196, 228, 271, 497, 500 ^V1^PNs for diazepam; n = 403, 375, 579, 590, 662 ^V1^PNs for PTZ, for 5, 10, 25, 50, and 100% respectively) and rTg4510 mice (right; n = 305, 299, 423, 495, 689 ^V1^PN for vehicle; n = 269, 208, 382, 430, 520 ^V1^PNs for diazepam; n = 322, 482, 407, 527, 503 ^V1^PN for PTZ, for 5, 10, 25, 50, and 100% respectively). (****) p<0.0001 by two-way ANOVA, genotype effect. Data are shown as mean ± SEM.

In the basal state, CC of excited ^V1^PN pairs was higher for rTg4510 mice compared to controls in the vehicle group (p<0.0001, by one-way ANOVA, genotype effect; Fig. 5B). Diazepam increased the CC of excited ^V1^PNs in both genotypes (p=0.0122, 0.0005 for controls and rTg4510, respectively, by one-way ANOVA, treatment effect; Fig. 5B) and a similar effect was observed with PTZ treatment (p<0.0001 for both genotypes, by one-way ANOVA, treatment effect).

In the evoked state, CC values of excited ^V1^PN pairs increased in a light intensity-dependent manner in vehicle-treated control mice (p<0.0001 by one-way ANOVA, intensity effect) but not in rTg4510 animals (Fig. 5C). Interestingly, diazepam restored this input-output relationship in rTg4510 mice (p<0.0001 between 5 and 100%, by one-way ANOVA, intensity effect), while disrupting it in the control group. On the other hand, PTZ disrupted the input-output pattern observed in vehicle-treated control mice, but not in rTg4510 mice.

### θ power is decreased in rTg4510 mice

As cortical pyramidal neurons activity pattern was altered in our early tauopathy model (Fig. 1-5), we performed EEG recordings from V1 to test whether differences in the basal and evoked states could also be detected in oscillatory activity of freely moving rTg4510 mice. Power spectral analysis of both basal and evoked activity are shown in Fig. 6A and 6B. Light stimulation did not significantly change the power spectrums in either genotype (Fig. 6). Quantification of the different frequency bands revealed a trend towards enhanced δ power in rTg4510 in the basal, but not evoked state, compared to controls, although it did not reach significance (p=0.2154, by one-way ANOVA, genotype effect; Fig. 6C). On the other hand, rTg4510 mice displayed a significantly reduced θ power compared to controls in both basal and evoked states (p<0.0001, by one-way ANOVA, genotype effect; Fig. 6D).

**Figure 6.**
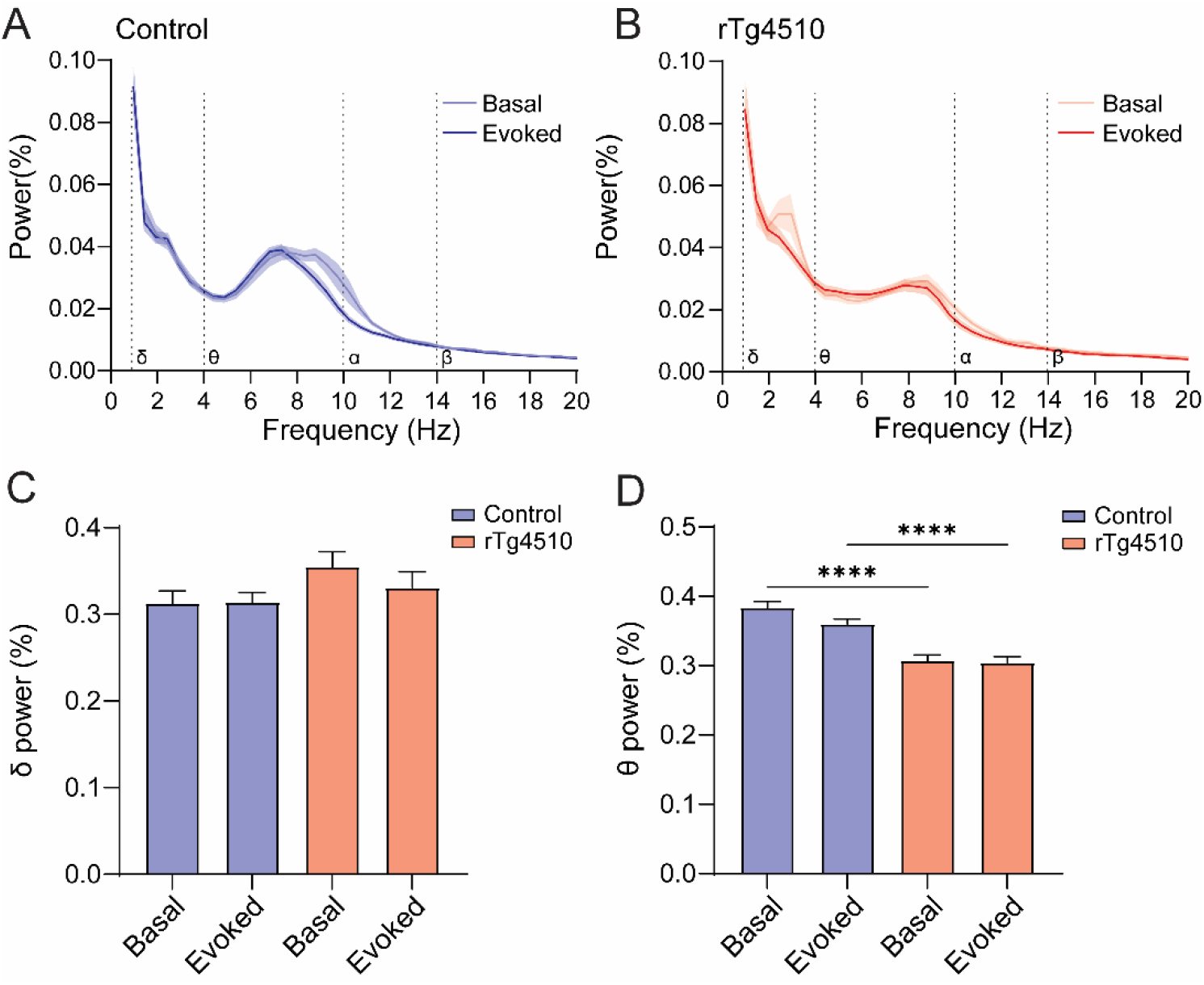
Altered EEG activity in rTg4510 mice. (A) EEG power spectra in control (blue, n=20) and (B) rTg4510 mice (red, n=17) in basal and evoked states. Shaded areas represent SEM. (C) δ and (D) θ power comparisons between control (blue, n=20) and rTg4510 mice (red, n= 17) in basal and evoked states. (****) p<0.0001 by two-way ANOVA, genotype and state effect. Data are shown as mean ± SEM.

### rTg4510 mice exhibit enhanced oscillatory and seizure-like activity in response to PTZ

It has been shown that tauopathy mouse models are more susceptible to epileptogenic treatments in the basal state [35, 45]. We therefore tested the effect of PTZ on EEG oscillations and epileptiform activity in freely moving control and rTg4510 mice. PTZ treatment did not change θ power in control animals, while it induced a marked increase in rTg4510 mice compared to baseline (p=0.003 for rTg4510, by one-way ANOVA, treatment; Fig. 7A, B). However, the absolute δ power was not different between rTg4510 and controls after PTZ treatment (Fig. S4). Next, EEG waveforms were analysed to test whether rTg4510 mice are more likely to display epileptiform activity triggered by the sub-convulsive dose of PTZ (Fig. 7C). Interestingly, rTg4510 showed greater susceptibility to PTZ with 60% of animals showing signs of short-lasting events of high amplitude oscillations within 8-10 Hz θ range whereas about 35% of control animals displayed absence-like seizures (responsive; Fig. 7D). The number of absence seizure episodes was significantly higher in responsive rTg4510 animals compared to controls (p=0.02, by student t-test; Fig. 7E). While the average duration of the episodes was not significantly altered between genotypes, a trend towards longer episode duration was observed in rTg4510 animals (p=0.0622 by student t-test; Fig. 7F). Finally, we found a significant correlation between the number of absence-like seizures and PTZ effect on θ activity only in responsive rTg4510 animals (p=0.012, by simple linear regression; Fig. 7G).

**Figure 7.**
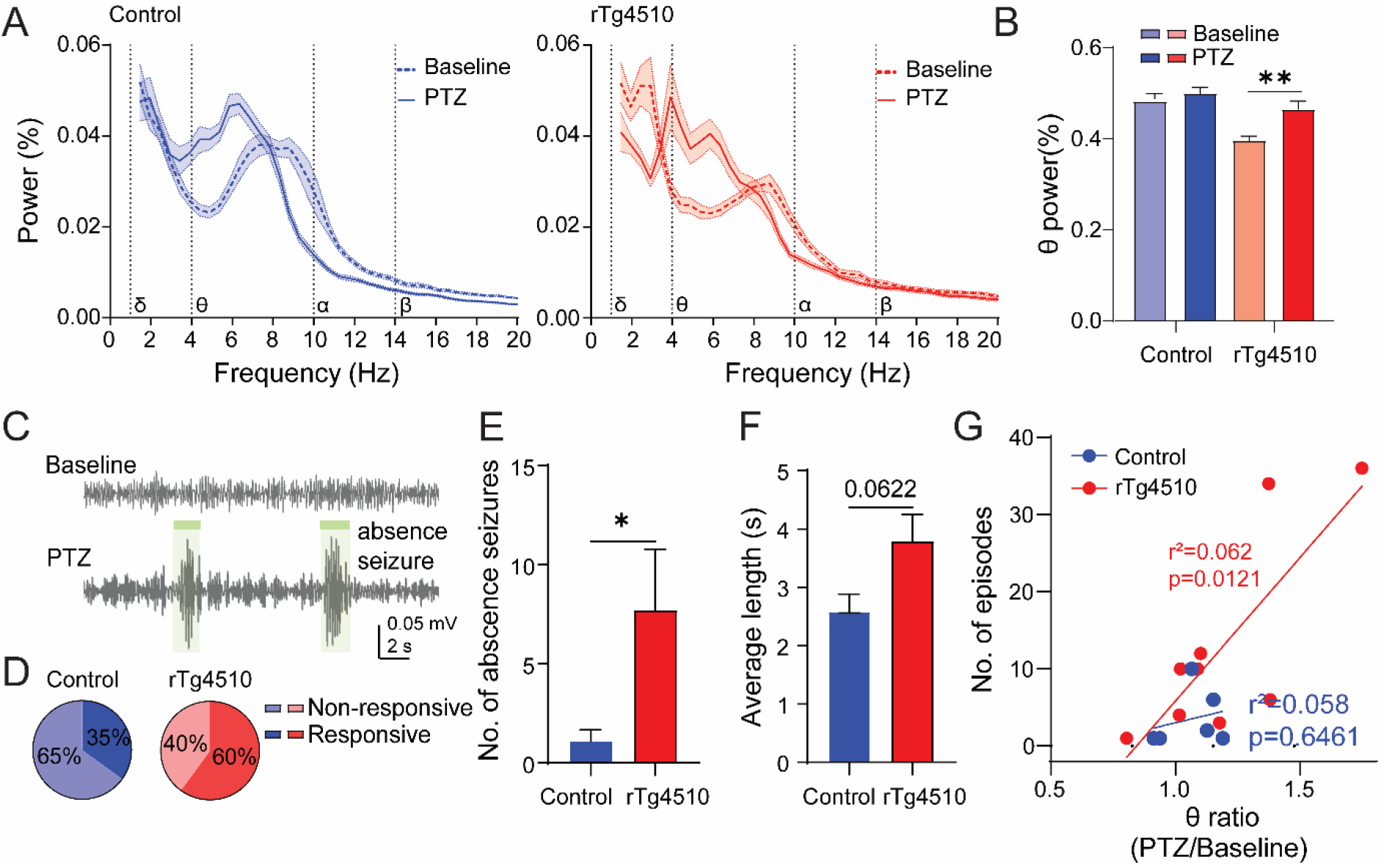
Abnormal PTZ effect on basal EEG oscillations in rTg4510 mice. (A) EEG power spectra of control (left; blue, n=20) and rTg4510 mice (right; red, n=17). (B) θ power comparisons between control (blue, n=20) and rTg4510 mice (red, n= 17) in baseline and after PTZ treatment. (**) p<0.01 by one-way ANOVA. (C) Exemplary EEG signature of absence seizures induced by PTZ. (D) Responsivity of control (blue, n=20) and rTg4510 mice (red, n=17) induced by PTZ treatment at a sub-threshold dose. An animal was considered responsive if it presented at least one absence seizure event. (E) Bar graphs showing the comparison of number of absence seizure events in control (blue, n=6) and rTg4510 animals (red, n=9) and (F) average length of the episode in control (blue, n=6) and rTg4510 mice (red, n=9). (*) p<0.05, by student t-test. (G) Correlation between the number of absence-like seizures and the effect of PTZ on θ power (ratio between θ power in PTZ and baseline). Data are shown as mean ± SEM.

## Discussion

Calcium imaging from thousands of genetically defined ^V1^PNs and visual cortical EEG recordings were employed to understand how the PN ensemble and oscillatory activity of the visual cortex is affected at early stages of tau pathology in transgenic mice prior to overt neurodegeneration. Our data provide a detailed functional profile of ^V1^PN impairments in basal and evoked states at the single cell level in a murine tauopathy model. We found an overall silencing of visual cortical pyramidal neurons in rTg4510 mice, which were insensitive to GABAergic pharmacological treatments. This was accompanied by aberrant increase in single cell synchrony in visual cortex which was, in some respects, restored by a sub-sedative dose of diazepam. Finally, EEG data demonstrate alterations in cortical oscillatory activity and increased susceptibility to epileptiform activity in tauopathy preceding neurodegeneration [9, 10]. Together with previously identified abnormal visual cortical responses in young rTg4510 animals [37], this study shows clear deficits in V1 pyramidal neurons ensemble excitability, synchrony and GABAergic tone which might all contribute to the abnormal oscillatory activity. These results provide a potential translational mechanism for understanding the neuronal basis of oscillatory activity impairments observed in AD [25, 46].

Visual evoked potentials were found to be altered in AD patients as well as in rTg4510 animals as young as 3 months old [37, 47]. While there is evidence for hippocampal and cortical PN function impairments in tauopathy mouse models, aberrant activity of pyramidal neurons of the primary visual cortex in basal and evoked states were never explored on a single cell level [5,14,34]. Miniature endoscope imaging in freely moving mice allowed us to evaluate whether neuronal activity pattern of ^V1^PNs was altered. First, we showed a strong decrease in ^V1^PNs input response to visual stimulation in rTg4510 animals compared to controls at the population and single-cell level. This effect was caused by a decrease in the fraction of excited neurons and an increase in the silenced ensemble, while no changes were observed in the general response parameters. Analogous to evoked state, a decline of ^V1^PNs activity in the basal state in rTg4510 mice compared to controls was observed. These findings are in agreement with independent reports showing silencing and attenuated excitability of hippocampal, entorhinal and parietal cortical neurons *in vitro*, *ex vivo* and *in vivo* in tauopathy models [20,29,48–50]. One hypothesis is that intrinsic ^V1^PN excitability might be impaired by distal relocation of axon initial segment resulting from tau hyperphosphorylation reported in hippocampal and cortical structures [34,50–52]. An alternative mechanism for the observed attenuation of cortical responsiveness could be related to an aberrant short- and/or long-range synaptic recruitment of ^V1^PNs (Fig 1, 2) [15,17,53,54]. In fact, hyperphosphorylated tau was shown to induce disruption of axonal transport necessary for effective synaptic transmission [53]. In addition, PN connectivity disruptions were reported in young rTg4510 and P301S mice in the form of reduction of hippocampal CA1 local field potential input-output slope upon Schaffer Collaterals stimulation and cortical layer V spine density reduction, respectively [17, 55].

As synchronicity amongst neurons together with oscillatory activity are also hallmarks of visual sensory representation [22,56,57], we further evaluated basal and evoked synchronous coactivation of ^V1^PNs at a single cell and population level using a cross-correlation analysis amongst neuronal pairs and EEG recordings in visual cortex V1 of freely moving rTg4510 mice, respectively [56, 57]. First, we observed a basal hyper-synchrony and impairment of input-related coactivation of ^V1^PN pairs in young rTg4510 animals (Fig. 5). As synchrony of ^V1^PN firing was shown to be fine-tuned by the visual cortical network to facilitate input detection and given that tau ablation altered synchronous activity in cortical neuronal cultures, our observations suggest aberrant single-cell visual temporal processing rigidity in rTg4510 mice [34,38,58]. Along with these findings, we discovered that oscillatory activity in the θ frequency was reduced in rTg4510 mice (Fig. 6). Interestingly, EEG oscillatory impairments were also observed in AD patients, who displayed reduction of power in 8-13 Hz frequency band [20,25,28]. As time-locked PN activity specifically resonates within the θ frequency [59], we hypothesize that the overall silencing of ^V1^PNs and their abnormal hyper-synchrony might impair fast θ rhythm driven by excitatory recurrent connectivity (Fig. 6) [60–62]. Our findings indicate that visual processing is not only impaired by a general decrease in signal magnitude, but that aberrant network synchronicity is another signature of early stages of tauopathy.

Network oscillations, visual response activity magnitude and synchrony amongst neurons are fine-tuned by local synaptic interaction with GABAergic interneurons [22,27,62–64]. Moreover, tau pathology might indirectly alter GABAergic transmission and consequently impair visual processing. For example, parvalbumin fast spiking neurons, known to control signal reliability of PNs in visual cortex and other brain areas, showed impairment in tauopathy and other AD mouse models [15,22,30,65– 70]. GABAergic pharmacological treatments were employed to evaluate whether visual cortex network processing aberration could be partially explained by alterations in GABA_A_R tone modulation of ^V1^PNs activity [64,71–73]. Our findings suggest a reduction of the GABAergic tone in rTg4510 mice. First, while diazepam and PTZ induced a respective decrease and increase in basal activity accompanied by an increase of silent ^V1^PNs neurons after diazepam administration in control mice, neuronal activity was insensitive to these compounds in rTg4510 animals (Fig. 3). Secondly, a sub-sedative dose of diazepam rescued the loss of correlation between visual stimulus and coactivation output of ^V1^PN neuronal pairs observed in rTg4510 mice (Fig. 5). Alongside our data, impairment in cortical GABA PET tracer uptake accompanied by reduction in GABAergic synaptic markers were demonstrated in 2 months old rTg4510 mice [15]. Interestingly, chronically administered sub-sedative doses of GABA_A_Rs benzodiazepine binding site ligands, such as diazepam and zolpidem, were shown to improve spatial learning/memory in the streptozotocin-induced AD rat model and rescue hippocampal long-term potentiation in a JNPL3 tauopathy mouse model by increasing synaptic density markers and soothing the neuroinflammation [72–74]. Although the clinical use of high doses of benzodiazepines revealed unsatisfactory results in the treatment of AD cognitive impairments, sub-sedative doses of diazepam (0.5-1 mg/kg) might have a beneficial effect [72,73,75]. Due to the widespread expression of GABA_A_Rs across various neural circuits, further studies targeting specific neuronal populations at a single cell level, such as specific GABAergic neurons, combined with network modelling will be crucial in strengthening the mechanistic understanding underlying the alteration of input detection observed in rTg4510 mice.

As spontaneous epileptiform activity in the frontal cortex and higher susceptibility to PTZ treatment was reported in another tauopathy mouse model, we used a sub-convulsive dose of PTZ to ascertain its influence on the oscillatory activity of the hyper-synchronous ^V1^PN network of rTg4510 mice [45,76,77]. PTZ enhanced single cell synchronicity and θ power was correlated with absence-like seizures in rTg4510 animals (Fig. 7). With pathological network synchrony and elevated θ power, rTg4510 mice proved to be more vulnerable to epileptiform activity during PTZ treatment compared to controls (Fig. 7) [78]. Thus, we propose that a more pronounced pathological hyper-synchronized network might result in higher susceptibility to epileptogenesis in rTg4510 mice. As hyper-synchronicity was displayed in epileptiform activity loci during seizures in patients and rats [79, 80] and tauopathy mouse models displayed higher susceptibility to epileptogenesis [35, 45], these findings point to a potential translatable mechanism underlying subclinical epilepsy that was often reported in AD patients [31, 32]. Building on the aforementioned studies, our data also suggest that tau pathology increases network vulnerability to epileptiform activity in absence of amyloid-β pathology, which is typically implicated in this pathophysiology [29, 81].

## Conclusion

Aberrant single-cell neuronal processing in visual cortex suggests that tau pathology at early stages might drive the observed EEG alterations. These findings provide a level of translatability to tauopathy in humans, as EEG is a frequently used non-invasive tool in clinical studies. Additionally, considering that drugs such a diazepam and zolpidem, have been found to rescue synaptic plasticity, cognition and neuroinflammation in tauopathy and other AD rodent models [73, 74], potential treatments aiming to positively modulate GABAergic action on neuronal circuits might act to suppress early network abnormalities and cognitive deficits in AD patients [75]. Overall, this study sheds light on neuronal activity alterations in rTg4510 animals and provides a new basis for untangling the complex mechanisms associated with tauopathy.

## Acknowledgements

We thank Elisa Corti (University of Coimbra, Portugal) for technical assistance on bulk analysis. This work was supported by the Innovation Fund Denmark (Ref. no. 9065-00019B).

## Supplementary material

**Table S1.**
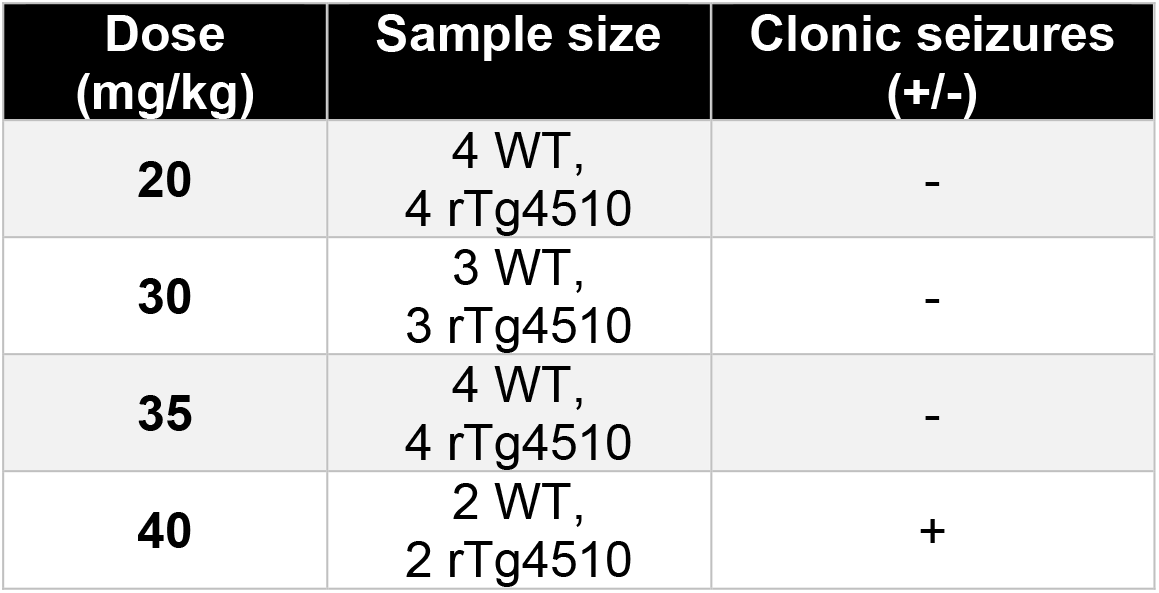
Pilot study investigating sub-convulsive doses of PTZ. If at least one animal from the test group developed myoclonic seizures, the dose was considered convulsive. WT – wild type mice.

**Figure S1.**
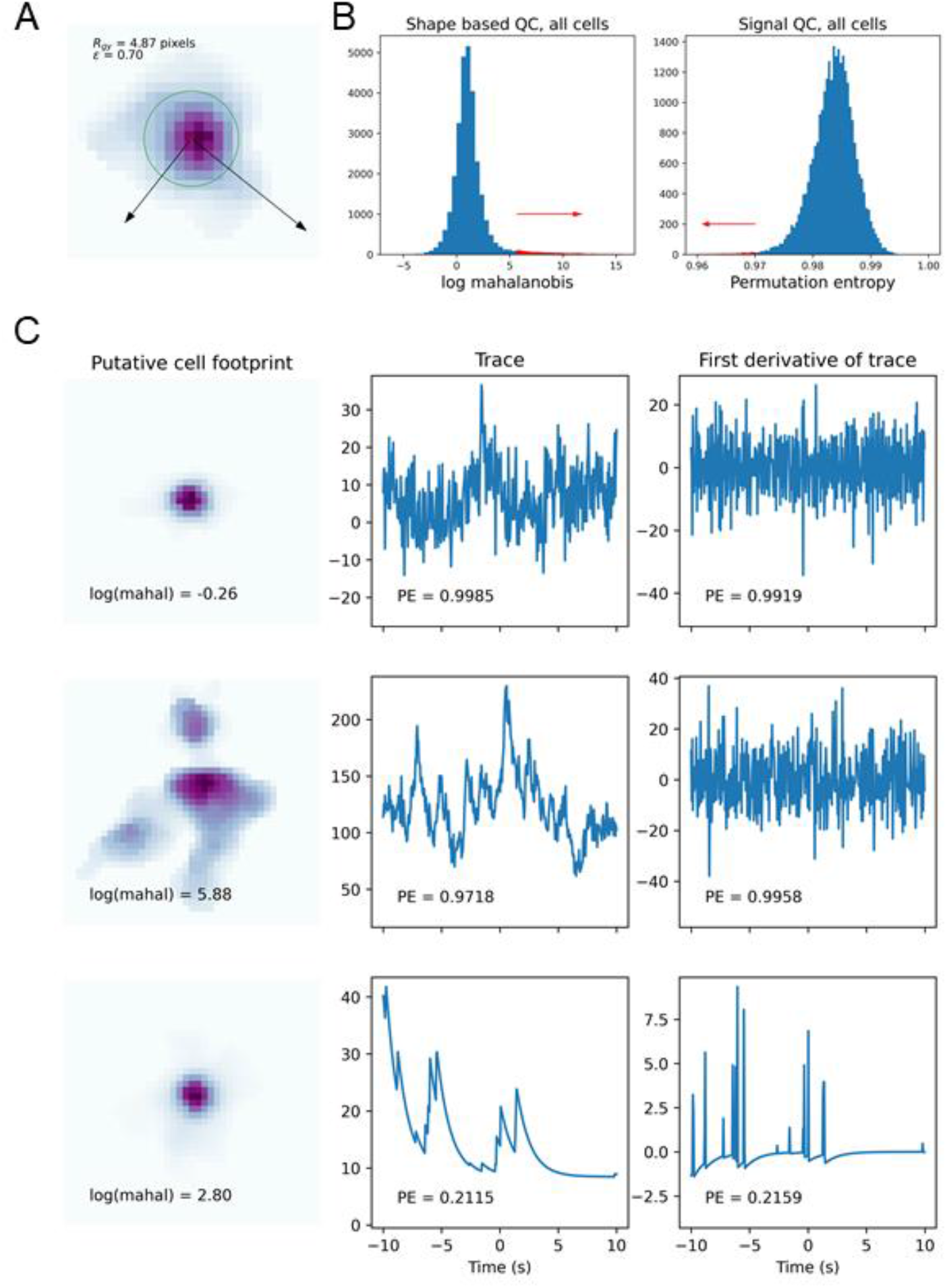
Illustration of the basic parameters for characterizing cell shape used in quality control (QC). (A) Image shows the pixel values of a putative cell. Red cross indicates the centroid if the cell image. The radius of gyration (𝑅_𝑔𝑦_) of the cell is represented as a green circle. The black arrows indicate the principal axes of the cell shape and their length illustrate the principal moments. The ratio of the principal moments (𝜖) is a measure of the elongation of the cell shape. (B) Histograms representing fraction of cells which passed (blue) or failed (red) QC in either shape (left) or signal (right) QC. (C) Three examples of cells and their scope in the QC measures. Top row shows footprint and trace of a cell that passes both shape and signal QC. The middle row shows a cell that fails in shape QC, but not signal QC, the lower row shows a cell that fails in signal QC. The log(mahal) and permutation entropy (PE) measures are shown in each figure.

**Figure S2.**
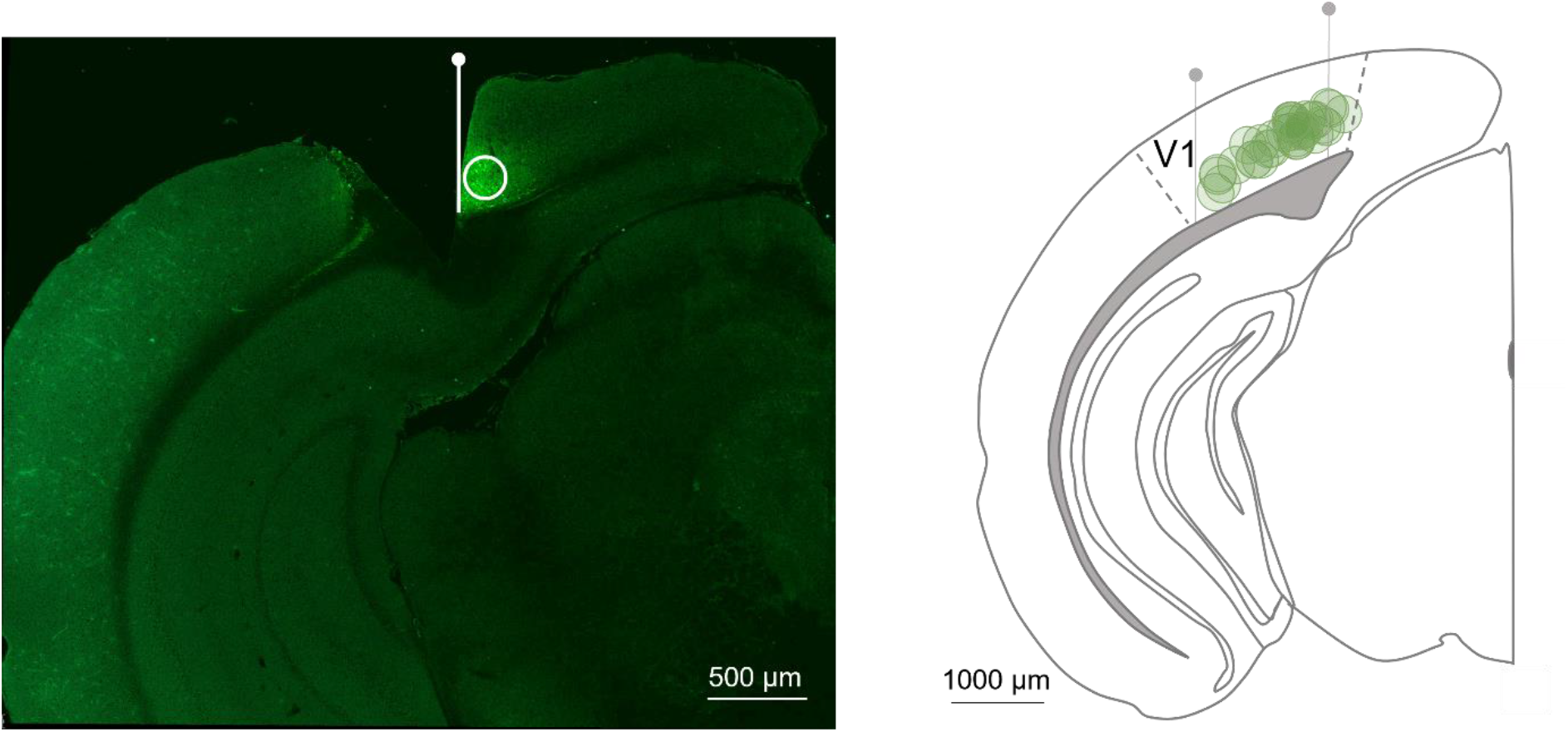
Approximate lens placement and virus expression. Left, representative image of successful lens placement and viral expression. Focal plane is represented by white line and centre of viral expression is labelled with a white circle. Right, qualitative representation of the most lateral and medial focal planes (grey lines) and centres of viral expression (green circles) from all imaged animals (n=18).

**Figure S3.**
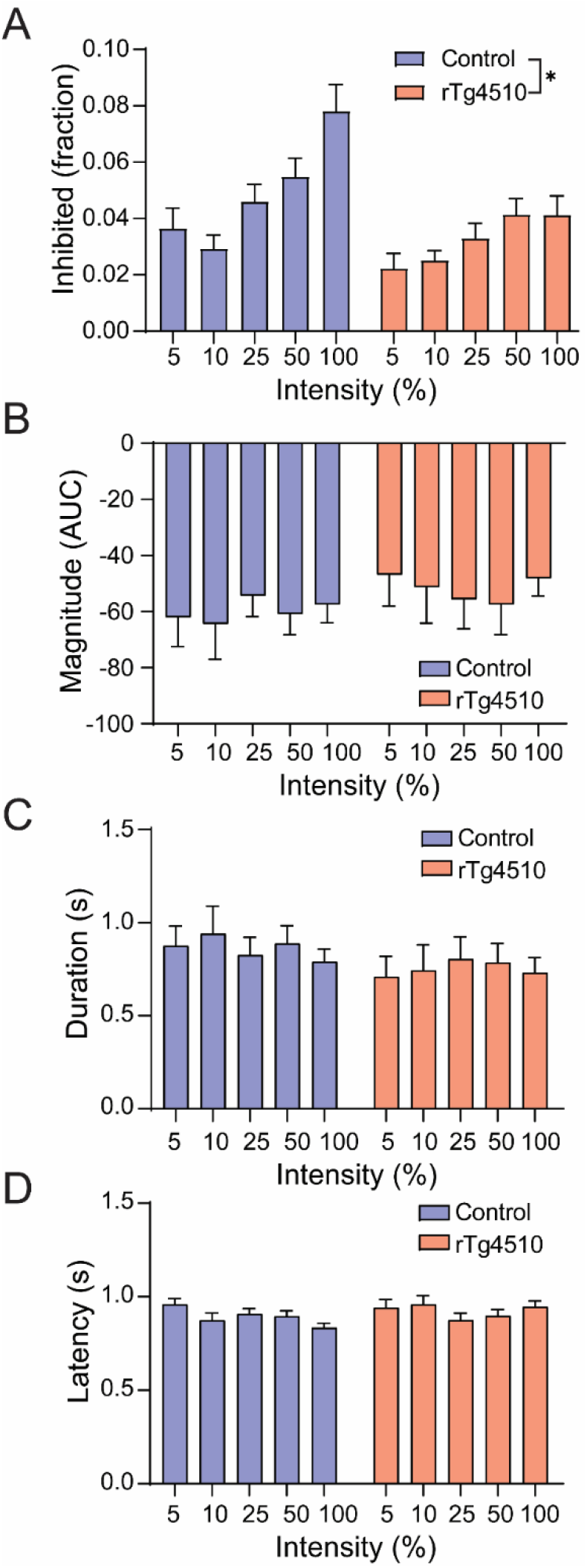
Decline in fraction of inhibited ^V1^PNs but no change in response parameters. (A) Inhibited ^V1^PNs fraction as a function of light intensity in control (blue, n=15) and rTg4510 mice (red, n=15). (*) p<0.05 by two-way ANOVA, genotype effect. (B) Magnitude, (C) duration and (D) latency of response of inhibited ^V1^PNs in control (blue, n=15) and rTg4510 mice (red, n=15). Data are shown as mean ± SEM.

**Figure S4.**
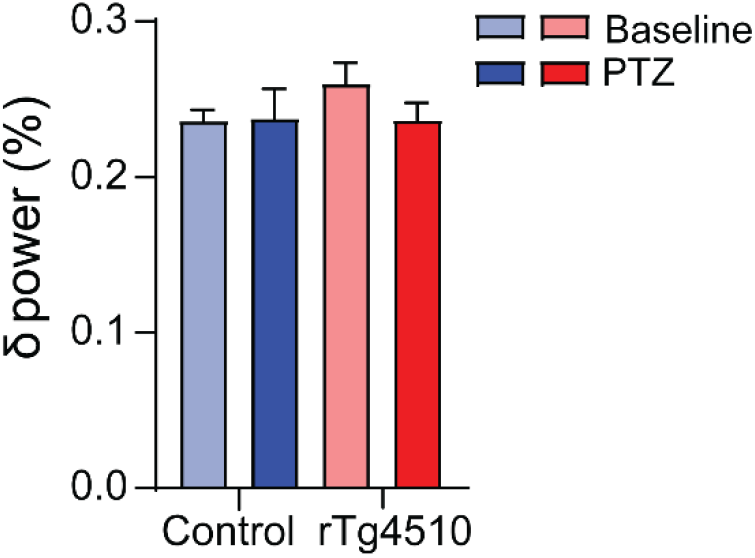
Total δ power in control (blue, n=20) and rTg4510 mice (red, n=17) pre- and post-PTZ administration. Data are shown as mean ± SEM.

## References

1. Arendt T, Stieler JT, Holzer M (2016) Tau and tauopathies. Brain Res Bull 126, 238–292.

2. Braak H, Braak E (1991) Neuropathological stageing of Alzheimer-related changes. Acta Neuropathol 82, 239–259.

3. Nilson AN, English KC, Gerson JE, Barton Whittle T, Nicolas Crain C, Xue J, Sengupta U, Castillo-Carranza DL, Zhang W, Gupta P, Kayed R (2016) Tau Oligomers Associate with Inflammation in the Brain and Retina of Tauopathy Mice and in Neurodegenerative Diseases. J Alzheimer’s Dis 55, 1083–1099.

4. Singh B, Covelo A, Martell-Martínez H, Nanclares C, Sherman MA, Okematti E, Meints J, Teravskis PJ, Gallardo C, Savonenko A V, Benneyworth MA, Lesné SE, Liao D, Araque A, Lee MK (2019) Tau is required for progressive synaptic and memory deficits in a transgenic mouse model of α-synucleinopathy. Acta Neuropathol 138, 551–574.

5. Rocher AB, Crimins JL, Amatrudo JM, Kinson MS, Todd-Brown MA, Lewis J, Luebke JI (2010) Structural and functional changes in tau mutant mice neurons are not linked to the presence of NFTs. Exp Neurol 223, 385–393.

6. Fox LM, William CM, Adamowicz DH, Pitstick R, Carlson GA, Spires-Jones TL, Hyman BT (2011) Soluble tau Species, Not Neurofibrillary Aggregates, Disrupt Neural System Integration in a tau Transgenic Model. J Neuropathol Exp Neurol 70, 588–595.

7. Jul P, Volbracht C, De Jong IEM, Helboe L, Brandt Elvang A, Pedersen JT (2016) Hyperactivity with Agitative-Like Behavior in a Mouse Tauopathy Model. J Alzheimer’s Dis 49, 783–795.

8. Blackmore T, Meftah S, Murray TK, Craig PJ, Blockeel A, Phillips K, Eastwood B, O’Neill MJ, Marston H, Ahmed Z, Gilmour G, Gastambide F (2017) Tracking progressive pathological and functional decline in the rTg4510 mouse model of tauopathy. Alzheimers Res Ther 9, 77.

9. SantaCruz K, Lewis J, Spires T, Paulson J, Kotilinek L, Ingelsson M, Guimaraes A, DeTure M, Ramsden M, McGowan E, Forster C, Yue M, Orne J, Janus C, Mariash A, Kuskowski M, Hyman B, Hutton M, Ashe KH (2005) Tau Suppression in a Neurodegenerative Mouse Model Improves Memory Function. Science (80-) 309, 476–481.

10. Helboe L, Egebjerg J, Barkholt P, Volbracht C (2017) Early depletion of CA1 neurons and late neurodegeneration in a mouse tauopathy model. Brain Res 1665, 22–35.

11. Mayford M, Bach ME, Huang Y-Y, Wang L, Hawkins RD, Kandel ER (1996) Control of Memory Formation Through Regulated Expression of a CaMKII Transgene. Science (80-) 274, 1678–1683.

12. Wang X, Zhang C, Szábo G, Sun Q-Q (2013) Distribution of CaMKIIα expression in the brain in vivo, studied by CaMKIIα-GFP mice. Brain Res 1518, 9–25.

13. Ramsden M, Kotilinek L, Forster C, Paulson J, McGowan E, SantaCruz K, Guimaraes A, Yue M, Lewis J, Carlson G, Hutton M, Ashe KH (2005) Age-Dependent Neurofibrillary Tangle Formation, Neuron Loss, and Memory Impairment in a Mouse Model of Human Tauopathy (P301L). J Neurosci 25, 10637–10647.

14. Crimins JL, Rocher AB, Luebke JI (2012) Electrophysiological changes precede morphological changes to frontal cortical pyramidal neurons in the rTg4510 mouse model of progressive tauopathy. Acta Neuropathol 124, 777– 795.

15. Shimojo M, Takuwa H, Takado Y, Tokunaga M, Tsukamoto S, Minatohara K, Ono M, Seki C, Maeda J, Urushihata T, Minamihisamatsu T, Aoki I, Kawamura K, Zhang M-R, Suhara T, Sahara N, Higuchi M (2020) Selective Disruption of Inhibitory Synapses Leading to Neuronal Hyperexcitability at an Early Stage of Tau Pathogenesis in a Mouse Model. J Neurosci 40, 3491–3501.

16. Kopeikina KJ, Polydoro M, Tai H-C, Yaeger E, Carlson GA, Pitstick R, Hyman BT, Spires-Jones TL (2013) Synaptic alterations in the rTg4510 mouse model of tauopathy. J Comp Neurol 521, 1334–1353.

17. Dalby NO, Volbracht C, Helboe L, Larsen PH, Jensen HS, Egebjerg J, Elvang AB (2014) Altered Function of Hippocampal CA1 Pyramidal Neurons in the rTg4510 Mouse Model of Tauopathy. J Alzheimer’s Dis 40, 429–442.

18. Crimins JL, Rocher AB, Peters A, Shultz P, Lewis J, Luebke JI (2011) Homeostatic responses by surviving cortical pyramidal cells in neurodegenerative tauopathy. Acta Neuropathol 122, 551–564.

19. Papanikolaou A, Rodrigues FR, Holeniewska J, Phillips KG, Saleem AB, Solomon SG (2022) Plasticity in visual cortex is disrupted in a mouse model of tauopathy. Commun Biol 5, 77.

20. Menkes-Caspi N, Yamin HG, Kellner V, Spires-Jones TL, Cohen D, Stern EA (2015) Pathological Tau Disrupts Ongoing Network Activity. Neuron 85, 959– 966.

21. Ishikawa A, Tokunaga M, Maeda J, Minamihisamatsu T, Shimojo M, Takuwa H, Ono M, Ni R, Hirano S, Kuwabara S, Zhang M-R, Aoki I, Suhara T, Higuchi M, Sahara N (2018) In Vivo Visualization of Tau Accumulation, Microglial Activation, and Brain Atrophy in a Mouse Model of Tauopathy rTg4510. J Alzheimer’s Dis 61, 1037–1052.

22. Zhu Y, Qiao W, Liu K, Zhong H, Yao H (2015) Control of response reliability by parvalbumin-expressing interneurons in visual cortex. Nat Commun 6, 1–11.

23. Booth CA, Witton J, Nowacki J, Tsaneva-Atanasova K, Jones MW, Randall AD, Brown JT (2016) Altered Intrinsic Pyramidal Neuron Properties and Pathway-Specific Synaptic Dysfunction Underlie Aberrant Hippocampal Network Function in a Mouse Model of Tauopathy. J Neurosci 36, 350–363.

24. Schön C, Hoffmann NA, Ochs SM, Burgold S, Filser S, Steinbach S, Seeliger MW, Arzberger T, Goedert M, Kretzschmar HA, Schmidt B, Herms J (2012) Long-Term In Vivo Imaging of Fibrillar Tau in the Retina of P301S Transgenic Mice. PLoS One 7, e53547.

25. Jeong J (2004) EEG dynamics in patients with Alzheimer’s disease. Clin Neurophysiol 115, 1490–1505.

26. Buzsaki G, Draghun A (2004) Neuronal oscillations in cortical networks. Science (80-) 304, 1926–1929.

27. Buzsáki G, Anastassiou CA, Koch C (2012) The origin of extracellular fields and currents — EEG, ECoG, LFP and spikes. Nat Rev Neurosci 13, 407–420.

28. Ahnaou A, Moechars D, Raeymaekers L, Biermans R, Manyakov N V, Bottelbergs A, Wintmolders C, Van Kolen K, Van De Casteele T, Kemp JA, Drinkenburg WH (2017) Emergence of early alterations in network oscillations and functional connectivity in a tau seeding mouse model of Alzheimer’s disease pathology. Sci Rep 7, 14189.

29. Busche MA, Wegmann S, Dujardin S, Commins C, Schiantarelli J, Klickstein N, Kamath T V, Carlson GA, Nelken I, Hyman BT (2019) Tau impairs neural circuits, dominating amyloid-β effects, in Alzheimer models in vivo. Nat Neurosci 22, 57–64.

30. Palop JJ, Mucke L (2016) Network abnormalities and interneuron dysfunction in Alzheimer disease. Nat Rev Neurosci 17, 777–792.

31. Scharfman HE (2012) “Untangling” Alzheimer’s Disease and Epilepsy. Epilepsy Curr 12, 178–183.

32. Scarmeas N, Honig LS, Choi H, Cantero J, Brandt J, Blacker D, Albert M, Amatniek JC, Marder K, Bell K, Hauser WA, Stern Y (2009) Seizures in Alzheimer Disease: Who, When, and How Common? Arch Neurol 66, 992– 997.

33. Friedman D, Honig LS, Scarmeas N (2012) Seizures and Epilepsy in Alzheimer’s Disease. CNS Neurosci Ther 18, 285–294.

34. Chang C-W, Evans MD, Yu X, Yu G-Q, Mucke L (2021) Tau reduction affects excitatory and inhibitory neurons differently, reduces excitation/inhibition ratios, and counteracts network hypersynchrony. Cell Rep 37, 109855.

35. Liu S, Shen Y, Shultz SR, Nguyen A, Hovens C, Adlard PA, Bush AI, Chan J, Kwan P, O’Brien TJ, Jones NC (2017) Accelerated kindling epileptogenesis in Tg4510 tau transgenic mice, but not in tau knockout mice. Epilepsia 58, e136– e141.

36. DeVos SL, Goncharoff DK, Chen G, Kebodeaux CS, Yamada K, Stewart FR, Schuler DR, Maloney SE, Wozniak DF, Rigo F, Bennett CF, Cirrito JR, Holtzman DM, Miller TM (2013) Antisense Reduction of Tau in Adult Mice Protects against Seizures. J Neurosci 33, 12887–12897.

37. Parka A, Volbracht C, Hall B, Bastlund JF, Nedergaard M, Laursen B, Botta P, Sotty F (2022) Visual evoked potentials as an early-stage biomarker in the rTg4510 tauopathy mouse model. bioRxiv.

38. Meeren HK., Van Luijtelaar ELJ., Coenen AM. (1998) Cortical and thalamic visual evoked potentials during sleep-wake states and spike-wave discharges in the rat. Electroencephalogr Clin Neurophysiol Potentials Sect 108, 306–319.

39. Yue M, Hanna A, Wilson J, Roder H, Janus C (2011) Sex difference in pathology and memory decline in rTg4510 mouse model of tauopathy. Neurobiol Aging 32, 590–603.

40. (2018) Inscopix Data Processing Software User Guide.

41. Pnevmatikakis EA, Soudry D, Gao Y, Machado TA, Merel J, Pfau D, Reardon T, Mu Y, Lacefield C, Yang W, Ahrens M, Bruno R, Jessell TM, Peterka DS, Yuste R, Paninski L (2016) Simultaneous Denoising, Deconvolution, and Demixing of Calcium Imaging Data. Neuron 89, 285–299.

42. Hubert M, Debruyne M, Rousseeuw PJ (2018) Minimum covariance determinant and extensions. Wiley Interdiscip Rev Comput Stat 10, e1421.

43. Bandt C, Pompe B (2002) Permutation Entropy: A Natural Complexity Measure for Time Series. Phys Rev Lett 88, 174102.

44. Friedrich J, Zhou P, Paninski L (2017) Fast online deconvolution of calcium imaging data. PLOS Comput Biol 13, e1005423.

45. García-Cabrero AM, Guerrero-López R, Giráldez BG, Llorens-Martín M, Ávila J, Serratosa JM, Sánchez MP (2013) Hyperexcitability and epileptic seizures in a model of frontotemporal dementia. Neurobiol Dis 58, 200–208.

46. Locatelli T, Cursi M, Liberati D, Franceschi M, Comi G (1998) EEG coherence in Alzheimer’s disease. Electroencephalogr Clin Neurophysiol 106, 229–237.

47. Moore NC (1997) Visual Evoked Responses in Alzheimer’s Disease: A Review. Clin Electroencephalogr 28, 137–142.

48. Angulo SL, Orman R, Neymotin SA, Liu L, Buitrago L, Cepeda-Prado E, Stefanov D, Lytton WW, Stewart M, Small SA, Duff KE, Moreno H (2017) Tau and amyloid-related pathologies in the entorhinal cortex have divergent effects in the hippocampal circuit. Neurobiol Dis 108, 261–276.

49. Hoover BR, Reed MN, Su J, Penrod RD, Kotilinek LA, Grant MK, Pitstick R, Carlson GA, Lanier LM, Yuan L-L, Ashe KH, Liao D (2010) Tau Mislocalization to Dendritic Spines Mediates Synaptic Dysfunction Independently of Neurodegeneration. Neuron 68, 1067–1081.

50. Hatch RJ, Wei Y, Xia D, Götz J (2017) Hyperphosphorylated tau causes reduced hippocampal CA1 excitability by relocating the axon initial segment. Acta Neuropathol 133, 717–730.

51. Druga R (2009) Neocortical Inhibitory System (cortical interneurons / GABAergic neurons / calcium-binding proteins / neuropeptides). Folia Biol 55, 201–217.

52. Grubb MS, Shu Y, Kuba H, Rasband MN, Wimmer VC, Bender KJ (2011) Short-and Long-Term Plasticity at the Axon Initial Segment. J Neurosci 31, 16049–16055.

53. Stamer K, Vogel R, Thies E, Mandelkow E-M, Mandelkow E-M (2002) Tau blocks traffic of organelles, neurofilaments, and APP vesicles in neurons and enhances oxidative stress. J Cell Biol 156, 1051–1063.

54. Hoffmann CA, Ochs NA, Burgold SM, Filser S (2012) Long-Term In Vivo Imaging of Fibrillar Tau in the Retina of P301S Transgenic Mice. PLoS One 7, 53547.

55. Hoffmann NA, Dorostkar MM, Blumenstock S, Goedert M, Herms J (2014) Impaired plasticity of cortical dendritic spines in P301S tau transgenic mice. Acta Neuropathol Commun 2, 1–11.

56. Babiloni C, Lizio R, Marzano N, Capotosto P, Soricelli A, Triggiani AI, Cordone S, Gesualdo L, Del Percio C (2016) Brain neural synchronization and functional coupling in Alzheimer’s disease as revealed by resting state EEG rhythms. Int J Psychophysiol 103, 88–102.

57. Pfurtscheller G, Lopes da Silva FH (1999) Event-related EEG/MEG synchronization and desynchronization: basic principles. Clin Neurophysiol 110, 1842–1857.

58. Minamisawa G, Funayama K, Matsumoto N, Matsuki N, Ikegaya Y (2017) Flashing Lights Induce Prolonged Distortions in Visual Cortical Responses and Visual Perception. eNeuro 4, 1–13.

59. Cardin JA, Carlén M, Meletis K, Knoblich U, Zhang F, Deisseroth K, Tsai LH, Moore CI (2009) Driving fast-spiking cells induces gamma rhythm and controls sensory responses. Nat 2009 4597247 459, 663–667.

60. Silva LR, Amitai Y, Connors BW (1991) Intrinsic Oscillations of Neocortex Generated by Layer 5 Pyramidal Neurons. New Ser 251, 432–435.

61. Budd JML (2005) Theta oscillations by synaptic excitation in a neocortical circuit model. Proc R Soc B Biol Sci 272, 101–109.

62. Cardin JA, Carlén M, Meletis K, Knoblich U, Zhang F, Deisseroth K, Tsai LH, Moore CI (2009) Driving fast-spiking cells induces gamma rhythm and controls sensory responses. Nature 459, 663–667.

63. Funayama K, Minamisawa G, Matsumoto N, Ban H, Chan AW, Matsuki N, Murphy TH, Ikegaya Y (2015) Neocortical Rebound Depolarization Enhances Visual Perception. PLOS Biol 13, e1002231.

64. Funayama K, Hagura N, Ban H, Ikegaya Y (2016) Functional Organization of Flash-Induced V1 Offline Reactivation. J Neurosci 36, 11727–11738.

65. Hampton DW, Webber DJ, Bilican B, Goedert M, Spillantini MG, Chandran S (2010) Cell-Mediated Neuroprotection in a Mouse Model of Human Tauopathy. J Neurosci 30, 9973–9983.

66. Verret L, Mann EO, Hang GB, Barth AMI, Cobos I, Ho K, Devidze N, Masliah E, Kreitzer AC, Mody I, Mucke L, Palop JJ (2012) Inhibitory interneuron deficit links altered network activity and cognitive dysfunction in Alzheimer model. Cell 149, 708–721.

67. Gonzalez-Burgos G, Cho RY, Lewis DA (2015) Alterations in Cortical Network Oscillations and Parvalbumin Neurons in Schizophrenia. Biol Psychiatry 77, 1031–1040.

68. Lovett-Barron M, Losonczy A (2014) Behavioral consequences of GABAergic neuronal diversity. Curr Opin Neurobiol 26, 27–33.

69. Jang HJ, Chung H, Rowland JM, Richards BA, Kohl MM, Kwag J (2020) Distinct roles of parvalbumin and somatostatin interneurons in gating the synchronization of spike times in the neocortex. Sci Adv 6, 5333–5355.

70. Petersen CCH (2014) Cell-type specific function of GABAergic neurons in layers 2 and 3 of mouse barrel cortex. Curr Opin Neurobiol 26, 1–6.

71. Kreuzer M, García PS, Brucklacher-Waldert V, Claassen R, Schneider G, Antkowiak B, Drexler B (2019) Diazepam and ethanol differently modulate neuronal activity in organotypic cortical cultures. BMC Neurosci 20, 58.

72. Pilipenko V, Narbute K, Pupure J, Rumaks J, Jansone B, Klusa V (2019) Neuroprotective action of diazepam at very low and moderate doses in Alzheimer’s disease model rats. Neuropharmacology 144, 319–326.

73. Levenga J, Krishnamurthy P, Rajamohamedsait H, Wong H, Franke TF, Cain P, Sigurdsson EM, Hoeffer CA (2013) Tau pathology induces loss of GABAergic interneurons leading to altered synaptic plasticity and behavioral impairments. Acta Neuropathol Commun 1, 34.

74. Umeda T, Kimura T, Yoshida K, Takao K, Fujita Y, Matsuyama S, Sakai A, Yamashita M, Yamashita Y, Ohnishi K, Suzuki M, Takuma H, Miyakawa T, Takashima A, Morita T, Mori H, Tomiyama T (2017) Mutation-induced loss of APP function causes GABAergic depletion in recessive familial Alzheimer’s disease: analysis of Osaka mutation-knockin mice. Acta Neuropathol Commun 5, 59.

75. Solas M, Puerta E, Ramirez M (2015) Treatment Options in Alzheimeŕs Disease: The GABA Story. Curr Pharm Des 21, 4960–4971.

76. Vossel KA, Ranasinghe KG, Beagle AJ, Mizuiri D, Honma SM, Dowling AF, Darwish SM, Van Berlo V, Barnes DE, Mantle M, Karydas AM, Coppola G, Roberson ED, Miller BL, Garcia PA, Kirsch HE, Mucke L, Nagarajan SS (2016) Incidence and impact of subclinical epileptiform activity in Alzheimer’s disease. Ann Neurol 80, 858–870.

77. Lam AD, Sarkis RA, Pellerin KR, Jing J, Dworetzky BA, Hoch DB, Jacobs CS, Lee JW, Weisholtz DS, Zepeda R, Westover MB, Cole AJ, Cash SS (2020) Association of epileptiform abnormalities and seizures in Alzheimer disease. Neurology 95, e2259–e2270.

78. Van Erum J, Dam D Van, De Deyn PP (2019) PTZ-induced seizures in mice require a revised Racine scale. Epilepsy Behav 95, 51–55.

79. Ren X, Brodovskaya A, Hudson JL, Kapur J (2021) Connectivity and Neuronal Synchrony during Seizures. J Neurosci 41, 7623–7635.

80. Dominguez LG, Wennberg RA, Gaetz W, Cheyne D, Snead OC, Perez Velazquez JL (2005) Enhanced Synchrony in Epileptiform Activity? Local versus Distant Phase Synchronization in Generalized Seizures. J Neurosci 25, 8077–8084.

81. Bezzina C, Verret L, Juan C, Remaud J, Halley H, Rampon C, Dahan L (2015) Early Onset of Hypersynchronous Network Activity and Expression of a Marker of Chronic Seizures in the Tg2576 Mouse Model of Alzheimer’s Disease. PLoS One 10, e0119910.

