## Supplementary material for "Early impairments of visually-driven neuronal ensemble dynamics in the rTg4510 tauopathy mouse model"

| Dose (mg/kg) | Sample size | Clonic seizures (+/-) |
|---|---|---|
| **20** | 4 WT,<br>4 rTg4510 | - |
| **30** | 3 WT,<br>3 rTg4510 | - |
| **35** | 4 WT,<br>4 rTg4510 | - |
| **40** | 2 WT,<br>2 rTg4510 | + |

Table S1. Pilot study investigating sub-convulsive doses of PTZ. If at least one animal from the test group developed myoclonic seizures, the dose was considered convulsive. WT – wild type mice.

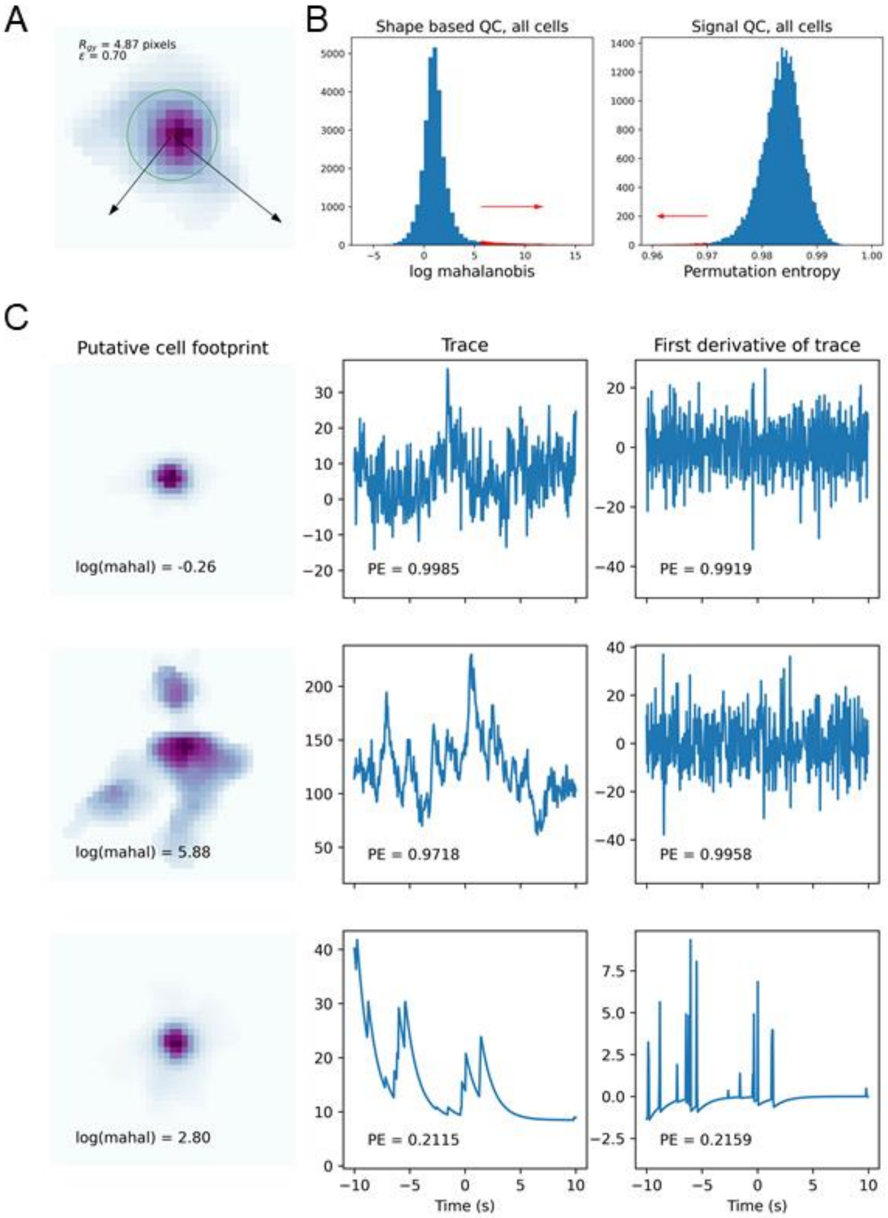


Figure S1. Illustration of the basic parameters for characterizing cell shape used in quality control (QC). (A) Image shows the pixel values of a putative cell. Red cross indicates the centroid if the cell image. The radius of gyration ($R_{gy}$) of the cell is represented as a green circle. The black arrows indicate the principal axes of the cell shape and their length illustrate the principal moments. The ratio of the principal moments ($\epsilon$) is a measure of the elongation of the cell shape. (B) Histograms representing fraction of cells which passed (blue) or failed (red) QC in either shape (left) or signal (right) QC. (C) Three examples of cells and their scope in the QC measures. Top row shows footprint and trace of a cell that passes both shape and signal QC. The middle row shows a cell that fails in shape QC, but not signal QC, the lower row shows a cell that fails in signal QC. The log(mahal) and permutation entropy (PE) measures are shown in each figure.

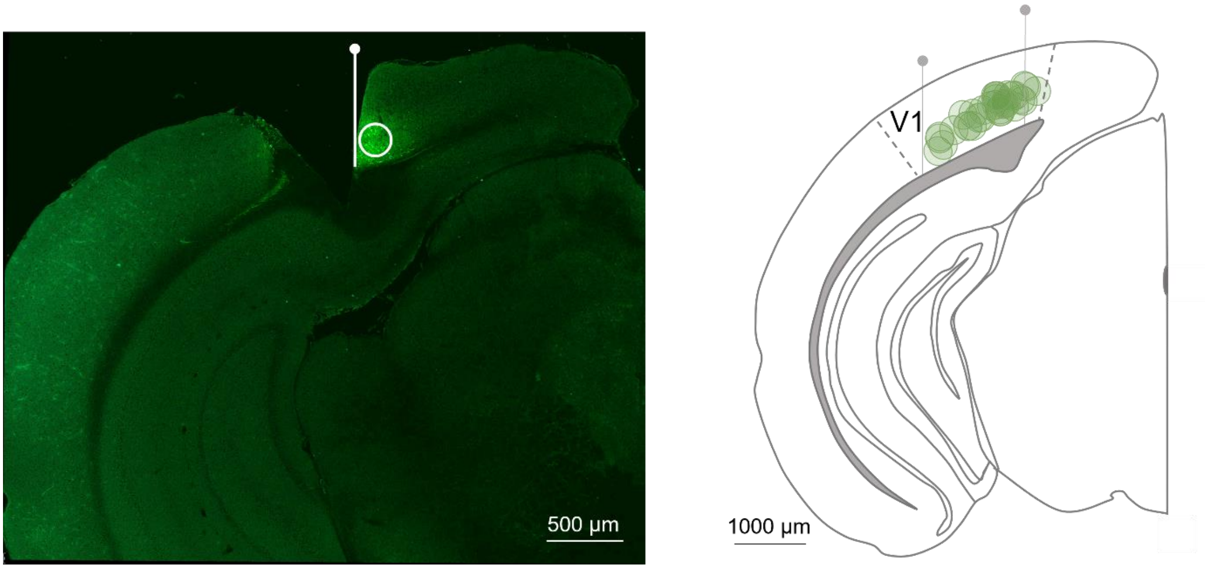


Figure S2. Approximate lens placement and virus expression. Left, representative image of successful lens placement and viral expression. Focal plane is represented by white line and centre of viral expression is labelled with a white circle. Right, qualitative representation of the most lateral and medial focal planes (grey lines) and centres of viral expression (green circles) from all imaged animals (n=18).

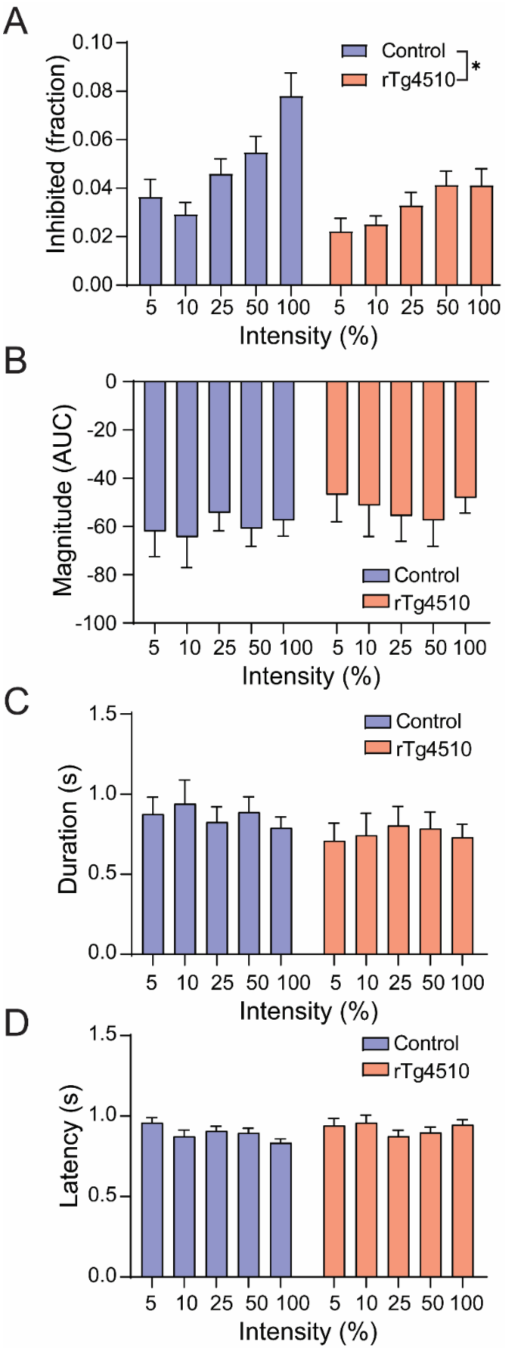


Figure S3. Decline in fraction of inhibited $^{V1}$PNs but no change in response parameters. (A) Inhibited $^{V1}$PNs fraction as a function of light intensity in control (blue, n=15) and rTg4510 mice (red, n=15). (*) $p<0.05$ by two-way ANOVA, genotype effect. (B) Magnitude, (C) duration and (D) latency of response of inhibited $^{V1}$PNs in control (blue, n=15) and rTg4510 mice (red, n=15). Data are shown as mean ± SEM.

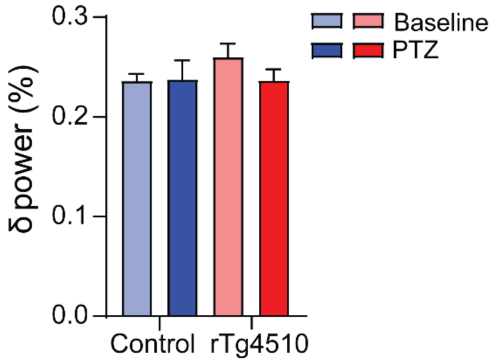


Figure S4. Total δ power in control (blue, n=20) and rTg4510 mice (red, n=17) pre- and post-PTZ administration. Data are shown as mean ± SEM.
